# A naturally arising broad and potent CD4-binding site antibody with low somatic mutation

**DOI:** 10.1101/2022.03.16.484662

**Authors:** Christopher O. Barnes, Till Schoofs, Priyanthi N.P. Gnanapragasam, Jovana Golijanin, Kathryn E. Huey-Tubman, Henning Gruell, Philipp Schommers, Nina Suh-Toma, Yu Erica Lee, Julio C. Cetrulo Lorenzi, Alicja Piechocka-Trocha, Johannes F. Scheid, Anthony P. West, Bruce D. Walker, Michael S. Seaman, Florian Klein, Michel C. Nussenzweig, Pamela J. Bjorkman

**Author notes:** These authors contributed equally (Christopher O. Barnes, Till Schoofs). Department of Biology, Stanford University, Stanford, CA, 94305, USA. GSK Vaccines, 1330 Rixensart, Belgium. Moderna Therapeutics, 200 Tech Square, Cambridge, MA, 02142, USA. Correspondence should be addressed to: Michel C. Nussenzweig and Pamela J. Bjorkman.

## Abstract

The induction of broadly neutralizing antibodies (bNAbs) is a potential strategy for a vaccine against HIV-1. However, most bNAbs exhibit features such as unusually high somatic hypermutation, including insertions and deletions, which make their induction challenging. VRC01-class bNAbs exhibit extraordinary breadth and potency, but also rank among the most highly somatically-mutated bNAbs. Here we describe a VRC01-class antibody isolated from a viremic controller, BG24, that has less than half the mutations of most other relatives of its class, while achieving comparable breadth and potency. A 3.8 Å X-ray crystal structure of a BG24-BG505 Env trimer complex revealed conserved contacts at the gp120 interface characteristic of the VRC01-class Abs, despite lacking common CDR3 sequence motifs. The existence of moderately-mutated CD4-binding site (CD4bs) bNAbs such as BG24 provides a simpler blueprint for CD4bs antibody induction by a vaccine, raising the prospect that such an induction might be feasible with a germline-targeting approach.

**Teaser:** An anti-HIV-1 antibody with comparable neutralization breadth and potency to similarly-classed antibodies, with half as many mutations.

## Introduction

In the last decade, it was discovered that a subset of HIV-1–infected individuals produce potent and broadly neutralizing antibodies (bNAbs) that target the HIV-1 envelope protein (Env) (*1–10*), a trimeric spike of gp120-gp41 heterodimers on the viral surface. Potent bNAb epitopes have been mapped across the entire surface of Env (*11–16*), have been shown to protect against and suppress infection in animal models (*17–20*), and exhibit antiviral activity in human clinical trials against circulating viral clades (*21–24*). Thus, it has been hypothesized that bNAbs could provide protection from HIV-1 infection if an efficient means of eliciting such antibodies could be developed (*25, 26*). However, while potent autologous neutralizing antibodies and neutralizing antibodies with intermediate breadth have been induced in wild type animal models (*27–30*), immunization strategies have yet to elicit potent, heterologous neutralizing bNAbs. Unusual features of HIV-1 bNAbs are considered one of the main barriers to their induction (*31*). Such features include high levels of somatic mutation, long heavy chain complementarity determining region 3 (CDRH3) loops, and insertions or deletions in antibody variable regions, all of which are rare features in the human repertoire and do not usually emerge until one to three years post-infection (*7, 32*).

Historically, CD4-binding site bNAbs are among the most broad and potent bNAbs (*33–35*). Members of the VRC01-class of CD4-binding site bNAbs, characterized by IGVH1-2*02 variable heavy (VH) gene segment use and an unusually short five-residue CDRL3 loop, have been isolated from many different donors and revealed to bind with a very similar orientation at the gp120 interface (*36*). However, these bNAbs are among the most heavily somatically mutated, creating a large barrier for induction of these antibodies in immunization strategies. Rational design has identified a minimally mutated version of VRC01-class antibodies that retains significant neutralization breadth and potency (*37*). However, it has been unclear whether such minimally mutated VRC01-class antibodies could arise naturally. Recently, a first example of a naturally-arising VRC01-class bNAb with a low mutation rate was described (*38*). The PCIN63 lineage showed similar features to VRC01-class bNAbs despite only 12-15% nucleotide somatic mutation compared to the putative germline V genes, providing evidence to a faster maturation route for VRC01-class bNAbs, where binding to the N276_gp120_-glycan may be an important first step and should be a consideration in VRC01-class priming immunogens.

Here, we describe antibody BG24, a VRC01-class bNAb, that targets the CD4-binding site with comparable neutralization breadth and potency to VRC01, while exhibiting half as many somatic mutations. We report CDR3 sequence motifs utilized by the BG24 lineage that are uncommon among VRC01-class bNAbs, challenging the notion of signature residues necessary for broad and potent neutralization. A 3.8Å crystal structure of BG24 Fab bound to a fully- and natively-glycosylated BG505 SOSIP.664 Env trimer (*39*) revealed a binding orientation consistent with VRC01-class bNAbs and contacts with an adjacent gp120 protomer. Collectively, these data provided the framework for engineering a minimally-mutated BG24 construct, which maintained breadth of binding to the mature construct and has direct implications in current HIV-1 immunization strategies.

## Results

### A family of VRC01-class antibodies in donor 391370 isolated by BG505-sorting

Donor 391370 was first diagnosed with HIV-1 in 1990 and was followed as part of the HIV Controller Consortium from 2005-2008 (*40*). The subject’s plasma from 2008 was previously tested against an early HIV-1 pseudovirus panel, showing broad and potent neutralizing activity (Table S1) (*9, 41*). To determine the epitope-specificity of 391370’s serum neutralizing activity, neutralization fingerprinting was done using the f61 pseudovirus panel (*42*) on a purified IgG sample from 2007, which showed a VRC01-class neutralization fingerprint (Fig. 1A,B). Indeed, a direct comparison with purified IgG of Patient 3, the subject from whom 3BNC117 (*9, 43*) was isolated, confirmed a very similar neutralization profile in breadth, potency and fingerprinting (Fig. 1A,B).

**Fig. 1.**
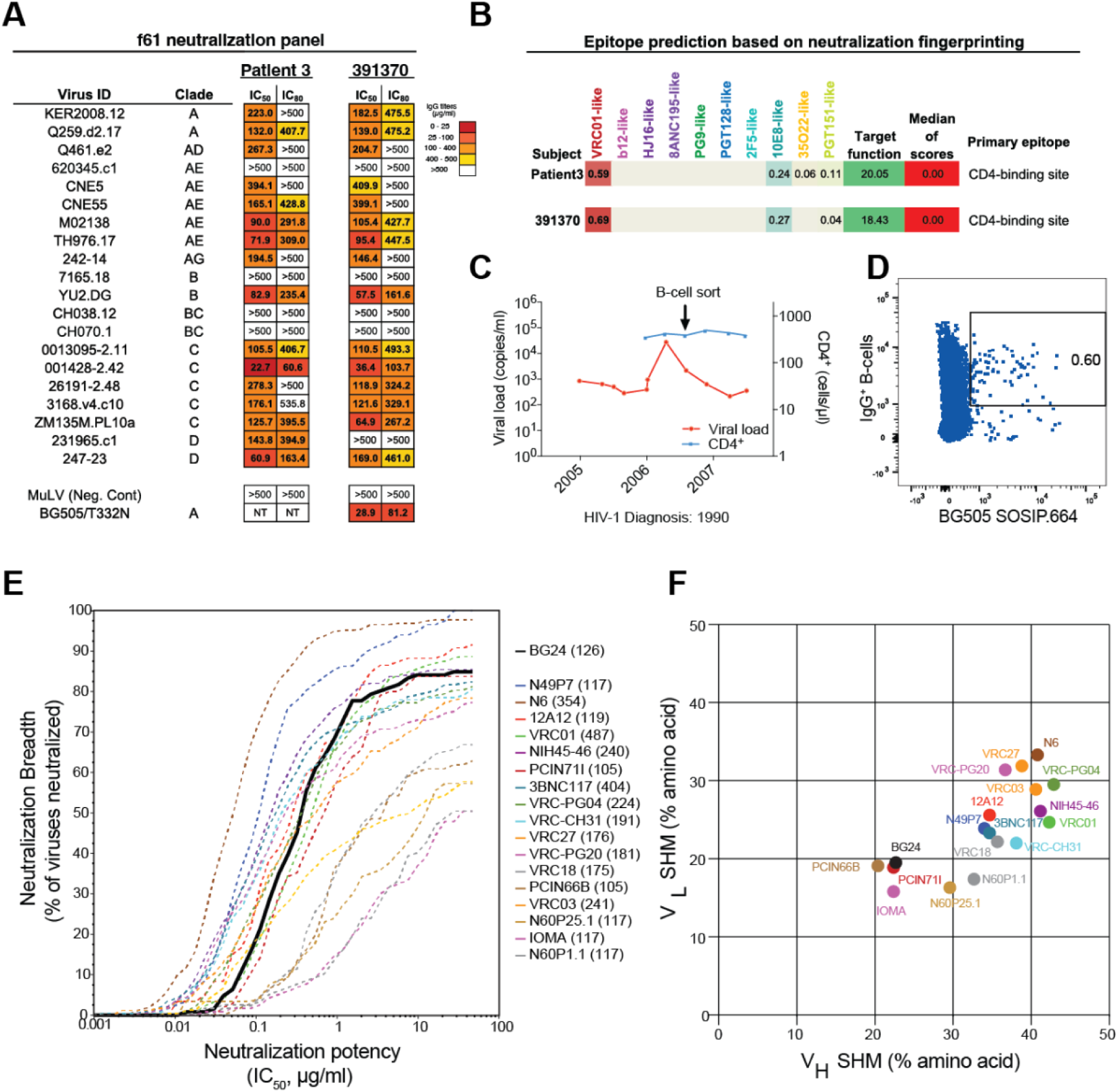
Isolation and characterization of antibody BG24 from donor 391370. A) Neutralization data of donor 391370’s serum IgG against a 20-virus fingerprinting panel (f61). The average median inhibitory concentrations (IC_50_) in µg/mL are shown from duplicate neutralization measurements. B) Fingerprinting analysis of f61 serum neutralization for Patient 3 and donor 391370. C) Plasma viral load and peripheral blood CD4^+^ T cell counts of donor 391370 over time. The arrow indicates the time point for BG505 SOSIP.664 bait-sorting. D) Sorting of single BG505 SOSIP.664^+^ IgG^+^ memory B-cells. E) Neutralization breadth and potency of BG24 on a 126-virus cross clade panel. Neutralization testing performed in duplicates, average shown. F) Somatic hypermutation (SHM) analysis of VRC01-like bNAbs, shown as % amino acid changes relative to germline variable gene sequence.

Due to strong neutralizing activity against BG505.T332N, single B cell sorting was carried out using BG505.SOSIP.664 (*39*) as a bait on a contemporaneous PBMC sample from 2007 (Fig. 1C,D) as described (*44*). From 20 million PBMCs, we recovered a total of 152 heavy chain and 159 light chain sequences from IgG^+^ memory B-cells (Fig. S1A). Both heavy and light chain sequences were highly clonal with 68% and 58% of sequences belonging to expanded groups of ≥ 2 clonally related members (Fig. S1A). Consistent with the VRC01-class fingerprint, the largest expanded clone was derived from an IGVH1-2*02 heavy chain germline gene segment. However, in contrast to the majority of VRC01-class antibodies, the members utilized a lambda and not a kappa light chain that was derived from the germline IGLV2-11*01 gene segment but showed the typical 5 amino acid length restriction in CDRL3 (Table S2).

Members of the IGVH1-2*02/IGLV2-11*01 clone showed quite a diverse phylogeny, but generally ranked lower in mutation count than other VRC01-class antibodies such as 3BNC117 and VRC01 (Table S2, Fig. S1B,C). Following production of monoclonal antibodies from 25 distinct members, we analyzed HIV-1 neutralizing activity against 5 viruses of the f61 panel that were best neutralized by 391370’s IgG (Fig. S1C). Clonal members exhibited a range of neutralization activity, and four members (BG5, BG24, BG33, BG38) with broad and potent anti-HIV activity were further tested on additional viruses of the f61 panel (Fig. S1D).

Clone member BG24 showed the most broad and potent neutralization activity, which recapitulated the serum neutralization profile of 391370’s IgGs with a strong CD4-binding site fingerprint (Fig. S1D). Consistent with this fingerprint, BG24 showed a mutational sensitivity profile similar to VRC01-class antibodies when tested against a HIV_YU2_ pseudovirus panel comprising escape mutations in common bNAb epitopes (Fig. S1E). Additional testing against the 12-virus global panel (*45*) showed BG24 to have comparable neutralization breadth to VRC01 and 3BNC117 (Fig. S1F). Moreover, BG24 exhibited equivalent breadth and potency to VRC01 against a 126-virus panel representative of all major circulating HIV-1 clades, neutralizing 85% of viruses with a geometric mean IC_50_ of 0.29 µg/ml (Fig. 1E, Table S3). As such, BG24 ranks among the most broad and potent of previously-described VRC01-class antibodies (Fig. 1E).

Surprisingly, BG24 showed one of the lowest numbers of somatic hypermutations of the clonal family, exhibiting 13.4% nucleotide (22.7% amino acid) and 8% nucleotide (19.5% amino acid) mutations in heavy and light chains, respectively. This mutation count is more than 2-fold lower than other VRC01-class antibodies (Fig. 1F, Fig. S2A, Table S2). In contrast to bNAbs 2F5 and 4E10 (*46*), no autoreactivity was found for BG24 by HEp-2 staining (Fig. S2B). To date, only one other patient-derived VRC01-class antibody (PCIN71I) with similar breadth, potency and low mutational count has been described (*38*). The discovery and characterization of BG24 and related modestly mutated CD4bs bNAbs suggests that targeting of this epitope may not require a high level of mutations to achieve breadth and potency.

### BG24 displays sequence features atypical of VRC01-class bNAbs

When aligned with other members of the VRC01-class bNAbs (Fig. S3A), BG24 shows conservation of sequence features such as R71_HC_, W50_HC_, N58_HC_, E96_LC_ and a deletion in CDRL1 to accommodate the gp120 N276-glycan (*36*). However, BG24 features a tyrosine at the -5 position in its CDRH3, a fixed position at the end of CDRH3 typically occupied by a tryptophan residue (*36*), and an uncommon 5-amino acid CDRL3 “SAFEY” sequence motif. Moreover, the presence of a potential N-linked glycosylation site (PNGS) at residue N58_HC_, which sits directly at the antigen-antibody interface, may alter BG24’s orientation at the CD4bs relative to other VRC01-class antibodies or reduce BG24’s ability to neutralize HIV-1 isolates.

Given that the Env binding orientations of VH1-2/VRC01-class bNAbs are highly convergent (*12, 33, 36, 47*), we speculated that glycosylation at position N58_HC_ of BG24 would likely reduce its potency and breadth. Previous studies of glycosylation patterns in eukaryotic proteins have shown that the presence of a N-x-S/T sequon is necessary but not sufficient for glycosylation, and moreover, that the N-x-S sequon is less frequently glycosylated than the N-x-T sequon (*48, 49*). Thus, to assess the impact of N58_HC_ glycosylation, we compared the neutralizing activity of a glycan-knockout construct (BG24 S60A_HC_) and a “glycan-occupied” construct (BG24 S60T_HC_) to the wild-type protein on the DeCamp global 12-strain panel (*45*). While we observed no significant difference in the neutralizing activity between the wild-type and BG24 S60A_HC_ construct, we observed an approximate 10-fold reduction in neutralization potency for the BG24 S60T_HC_ construct relative to wild-type (Fig. S3B). This result suggests that: i) the BG24 S60T_HC_ construct has a higher N-glycan occupancy at position N58_HC_ relative to wild-type BG24, and ii) glycosylation at position N58_HC_ is positively correlated with reduced BG24 neutralizing activity, which is likely explained by BG24 adopting a similar Env binding orientation as other VRC01-class bNAbs.

### Structure of BG24-Env complex shows similar recognition of gp120 as VRC01-class bNAbs

Since we observed no difference in the neutralization profile of BG24 S60A_HC_ compared to wild-type, we used the S60A_HC_ construct to define the Env binding mechanism of BG24 to eliminate potential interference of a glycosylated N58_HC_ residue in structural studies. We determined a 2.0 Å crystal structure of the BG24 S60A_HC_ Fab and a 3.8 Å crystal structure of BG24 S60A_HC_ Fab in complex with a natively-glycosylated clade A BG505 SOSIP.664 trimer and a Fab from the V3- glycan targeting bNAb 10-1074 (Table S4 and Fig. 2A). Comparison of the BG24 Fab components of the two structures revealed that BG24 did not undergo large conformational changes upon Env binding (Fig. S4A, root mean square deviation of 0.9 Å when aligned against 219 Cα atoms comprising the BG24 variable domains). Relative to VRC01-class bNAbs, BG24 maintained a similar gp120-binding orientation, consistent with an overall epitope focused on the portion of the CD4bs within the gp120 outer domain, framed by the N197_gp120_, N276_gp120_, and N363_gp120_ glycans (Figs. 2B and S4B). With the exception of CDRL2, all BG24 CDRs were involved in gp120 recognition, burying a similar degree of surface area on the CD4bs loop, D loop, and the V5-loop of gp120 as VRC01, VRC03 and 3BNC117 (Fig. S4C,D). Typical interactions between VRC01-class bNAbs and gp120 are conserved with BG24 including: i) a salt bridge between R71_HC_ and D368_gp120_, ii) potential hydrogen bonding between W50_HC_ and N280_gp120_, and iii) potential hydrogen bonding between N58_HC_ and the backbone carbonyl of R456_gp120_ (Fig. 2C).

**Fig. 2.**
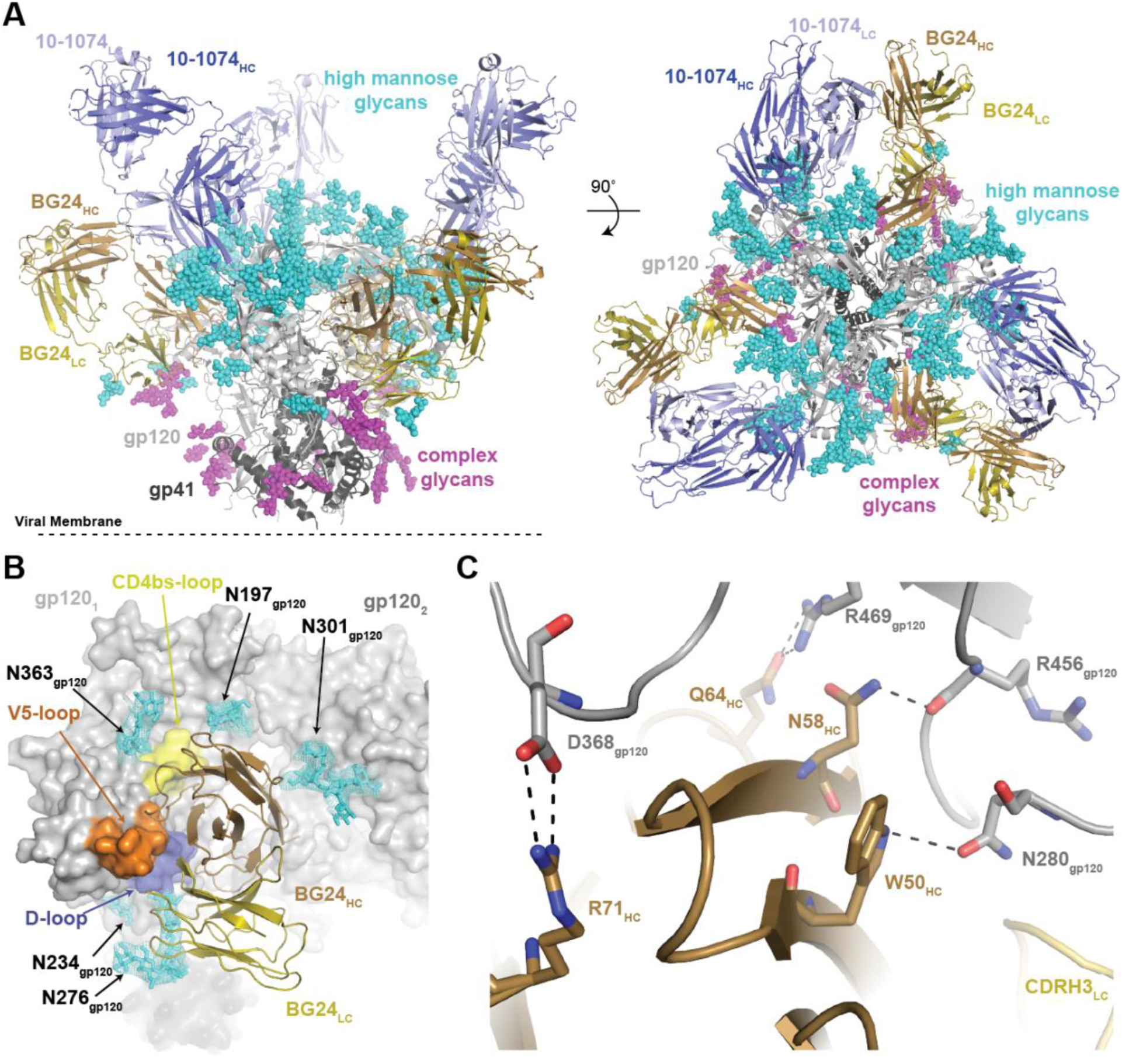
BG24 recognition of HIV-1 Env has features in common with VRC01-class bNAbs. A) Side and top views of the 3.9Å X-ray structure of the BG24-BG505-10-1074 complex colored by components (dark gray, gp41; light gray, gp120; shades of blue, 10-1074 Fab; shades of brown, BG24 Fab). B) Surface representation of gp120 (gray), with main loops at the CD4bs colored (yellow, CD4bs-loop; blue, D-loop; orange, V5-loop) and BG24 shown as cartoon representation. N-linked glycans modeled in the structure are shown as cyan sticks with electron density contoured at 1.5σ. C) Stick representation of residue level contacts between VRC01-class signature residues in BG24_HC_ (brown) with gp120 (gray). Dashed black lines indicated potential for H-bond interactions.

In addition to the characteristic VH1-2 contacts, the BG24 epitope includes interprotomer interactions that have been observed for other VRC01-class antibodies (Fig. S4C,D)(*50, 51*). However, unlike 3BNC117 or VRC03, which utilize insertions in HC FWR3 to contact the V3-loop base on the adjacent protomer (*50, 52*), BG24’s CDRH1 interacts with α0 residues of the adjacent gp120 protomer, with N28_HC_ potentially hydrogen bonding with the backbone carbonyl of E64_gp120_ (Fig. 4E). Moreover, BG24’s binding orientation brings HC FWR1 into close proximity with the neighboring N301_gp120_ glycan (modeled in the density as a complex-type biantennary N-glycan) burying ∼127 Å^2^ of glycan surface area (Fig. S5A,B). This interaction is not unique to BG24, having been observed in crystal structures of a VRC01-bound high-mannose fully-glycosylated Env trimer (*53*) and an IOMA-bound natively- and fully-glycosylated Env trimer (*12*). However, in contrast to previous studies that attributed a shift to more positively-charged antigen combining sites of VRC01-class bNAbs to interactions with complex-type N-glycans (*47*), comparison of the N301_gp120_ glycan binding surface on BG24 relative to germline VRC01 showed minimal changes in the electrostatic surface potential (Fig. S5C-E). This result suggests that unlike the complex-type N197_gp120_ and N276_gp120_ glycans that frame the CD4bs, the complex-type N301_gp120_ glycan plays little to no role in VRC01-class bNAb maturation.

### Non-traditional CDR3 sequence motifs favorably interact with HIV-1 gp120

In addition to low numbers of somatic mutations, BG24 is defined by CDR3 features that are uncommon among the VRC01-class antibodies. BG24 utilizes a 5-amino acid length CDRL3 with an unusual “SAFEY” sequence motif that differs from the consensus “QQYEF” motif of κ^+^ VRC01-class bNAbs and is distinct among λ^+^ VRC01-class bNAbs (Fig. S3A) (*36, 54*). Despite the distinct CDRL3 sequence motif, BG24 includes a negatively-charged Glu residue at LC position 96 that maintains signature H-bond interactions with N280_gp120_ and G459_gp120_ in the D- and V5-loops, respectively (Fig. 3A). Additionally, E96_LC_ potentially interacts with R456_gp120_, a contact not routinely found in VRC01-class antibody-Env structures and only previously observed in an IOMA-BG505 Env structure due to IOMA’s 8-residue CDRL3 (*12*).

**Fig. 3.**
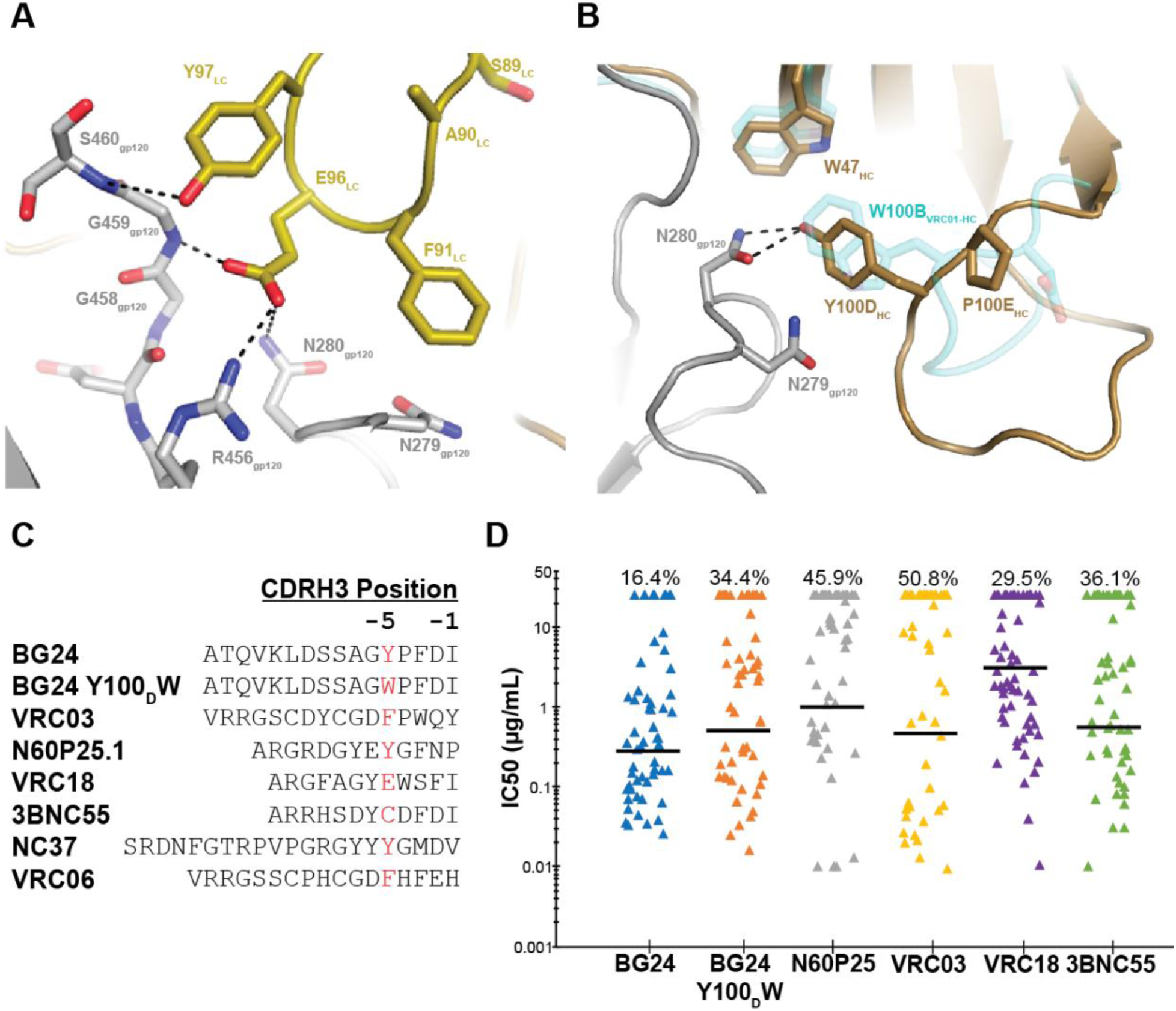
BG24’s CDR3 sequence motifs are uncommon among VRC01-like bNAbs. A) Stick representation of residue level contacts between residues in BG24’s CDRL3 loop (yellow) with gp120 (gray). Potential H-bond interactions are shown as black dashed lines. VRC01 CDRH3 is also shown in **panel B** (cyan). B) Stick representation of residue level contacts between residues in BG24’s CDRH3 loop (brown) with gp120 (gray). The CDRH3 loop of VRC01 (PDB 6VX8) is also shown (cyan). Potential H-bond interactions are shown as black dashed lines. C) CDRH3 sequence alignment of VRC01-like antibodies that lack a Trp residue at the -5 position. D) Neutralization data for in-common (n=61) cross-clade viruses of BG24 and VRC01-like antibodies that lack a Trp residue at the -5 position. The geometric mean IC_50_ value against antibody-sensitive strains is indicated by the horizontal black line. The percentage of non-neutralized strains is indicated on the top for each antibody. Analysis of neutralization and graphing was done using the Antibody Database (v 2.0) (*73*).

In the heavy chain, BG24 lacks the canonical Trp residue at the -5 position in its CDRH3, which is highly-conserved in most VRC01-class antibodies and forms H-bond interactions with residue N279_gp120_ at that antibody-antigen interface (*36*). Interestingly, enrichment of non-Trp residues at the -5 position of CDRH3 in naïve B-cells sorted with eOD-GT8, a VRC01-class targeting immunogen (*55, 56*), suggested that recombination of the CDRH3 with the low frequency IGHJ2*01 gene segment is not a requirement of VRC01-class B-cell precursors, and a Trp residue in this position could plausibly arise during affinity maturation (*54*). BG24 features a Tyr at this position, which preserves D-loop interactions at the interface by forming a H-bond with a side chain oxygen atom on N280_gp120_ (Fig. 3B). To test whether BG24 would show increased anti-HIV activity with a Trp residue in this position (Kabat numbering - CDRH3 residue 100D), we generated a Y100_D_W construct and assayed neutralizing activity against a 126 virus panel. We observed BG24 Y100_D_ to be broader and more potent than BG24 W100_D_, as well as all other VRC01-class bNAbs lacking a Trp residue at the -5 position in CDRH3 (Fig. 3C,D and Table S3). Thus, our results demonstrate the potential for potency and breadth of VRC01-class antibodies encoding for non-Trp residues at this position, which is frequently observed in naïve B-cells sorted with CD4bs immunogens (*54*).

### Substitutions in BG24 CDRH2 loop improve neutralizing activity

Previous studies have shown that VRC01-class antibody contacts with the gp120 inner domain (*57*), “Phe43 pocket” (*58–60*), or interprotomer interactions (*51, 52*) enhance antibody activity. Given BG24’s low number of mutations, we sought to enhance BG24’s potent neutralization and breadth by incorporating substitutions in CDRH2 residues that mimic Phe43 pocket filling. In addition to the G54W mutation that was engineered into the VRC01-class bNAbs NIH45-46 and VRC07 (*58, 60*), we designed BG24 constructs that substituted CDRH2 residues from VRC-PG20 (Fig. S6A), a IGVH1-2*02 VRC01-class antibody that utilizes a λ^+^ light chain (IGVL2-14) and encodes a W54 residue in CDRH2 (*61*). We postulated that residues flanking the large aromatic substitution at position 54 potentially reduce polyreactive recognition of non-HIV-1 antigens previously observed for the G54W substitution (*62*). To assess potential polyreactivity of BG24 and the BG24-derived constructs, we utilized a baculovirus-based polyreactivity assay (*63*). While antibodies NIH45-46^G54W^, 2F5 and 4E10, (which are known to be polyreactive) showed strong signals, no evidence of polyreactivity was found for BG24 or any of the designed constructs (Fig. S6B).

We next tested our constructs for neutralizing activity on the 12-strain global panel (*45*) and compared potency and breadth against BG24. In general, we observed a 2-5-fold improvement in IC_50_ values for all constructs relative to unmodified BG24, with constructs containing VRC-PG20 CDRH2 sequences being the most potent (Fig. 4A). When tested against a 126 virus panel, the engineered BG24 constructs achieved ∼90% breadth and a 2-3 fold improvement in the geometric mean IC_50_ value relative to BG24 (Table S3: BG24 – IC_50_=0.29µg/mL / 84.9% breadth; BG24_G54W_ – IC_50_=0.15µg/mL / 88.1% breadth; BG24_PG20-CDR2-v1_ – IC_50_=0.15µg/mL / 92.9% breadth; BG24_PG20-CDR2-v2_ – IC_50_=0.14µg/mL / 92.1% breadth).

**Fig. 4.**
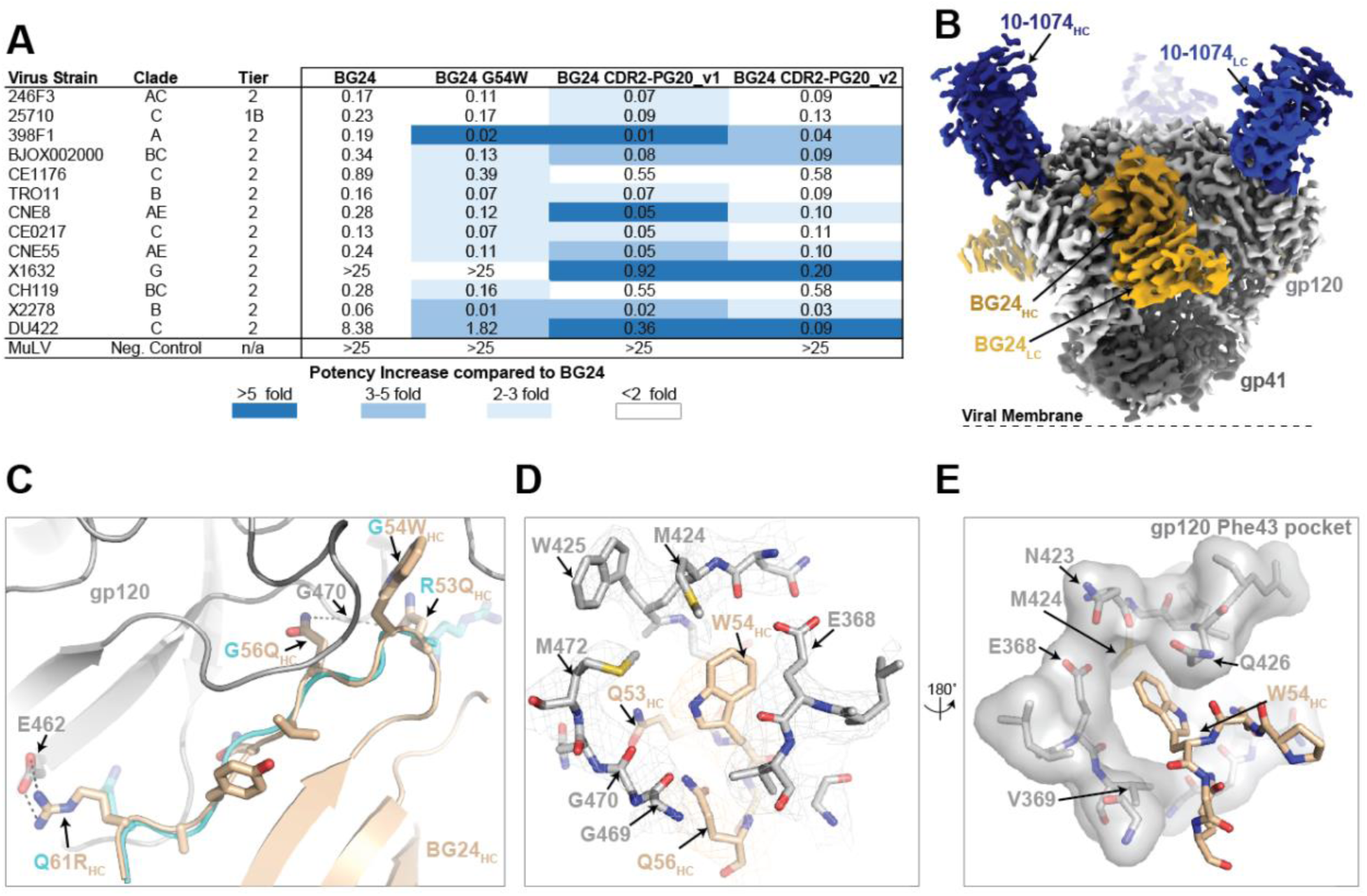
Improvements to BG24 neutralization potency and breadth. A) Neutralization data of engineered BG24 constructs against the global 12 virus panel. The average mean IC_50_ in µg/mL are shown from duplicate neutralization measurements. B) Side and top views of the 3.5Å single-particle cryo-EM reconstruction of the BG24_PG20-CDR2-v2_-DU422-10-1074 complex colored by components (dark gray, gp41; light gray, gp120; shades of blue, 10-1074 Fab; shades of brown, BG24 Fab) C) Cartoon and stick representation of BG24_PG20-CDR2-v2_ CDRH2 (wheat) at the gp120 (gray) interface. The CDRH2 loop from the BG24-BG505 complex (cyan) is overlaid with amino acid mutations between the two constructs labeled. Potential H-bond interactions are shown as black dashed lines. D) Modeling of the Phe43 gp120 pocket with BG24_G54W_ pocket-filling mutation highlighted (wheat). Electron density contoured at 7σ is shown for gp120 and BG24. E) Surface representation of gp120 Phe43 pocket (gray) and BG24_G54W_ pocket-filling mutation (wheat).

To understand the basis of this increased neutralizing activity, we solved a 3.5 Å single-particle cryo-EM structure of a natively-glycosylated DU422 SOSIP.664 v4.1 trimer in complex with BG24_PG20-CDR2-v2_ and 10-1074 Fabs (Fig.s 4B, S6C-F and Table S5). Consistent with previous observations(*58, 60*), the W54_HC_ residue encoded by CDRH2 is accommodated within gp120’s Phe43 pocket, increasing contacts with the gp120 inner domain (Fig. 4C-E). Additionally, N53_HC_ and N56_HC_ form backbone potential backbone interactions with G469-G470_gp120_ and R61_HC_ establishes an additional salt bridge with E462_gp120_, interactions not observed with the parent BG24 antibody. Collectively, these interactions add an additional ∼170 Å^2^ of buried surface area on the antibody paratope by providing favorable interactions that likely increase BG24’s affinity to the CD4bs epitope, explaining the enhanced neutralization activity of against DU422.

### BG24 has comparable *in vivo* efficacy to VRC01

Anti-HIV-1 bNAbs are being considered as agents for HIV-1 treatment and prevention. A few antibodies have already been tested in human subjects, and been found to have therapeutic activity, including antibody VRC01 (*23, 64, 65*). To determine whether BG24 shows therapeutic potential *in vivo*, we sought to compare its anti-HIV activity with well-established antibody VRC01 in HIV_YU2_-infected humanized mice. While the pharmacokinetics of VRC01 are known, we first evaluated the pharmacokinetic properties of BG24 by intravenous injection into non-humanized NOD-Rag1^null^ IL2rg^null^ (NRG) mice (n=6). BG24 showed a similar decline in serum to other VRC01-class antibodies indicating an acceptable pharmacokinetic profile (Fig. S7) (*34*).

We then infected humanized NRG mice intraperitoneally with HIV_YU-2_ and subsequently treated them subcutaneously with repeated monotherapy of antibody BG24 (n=6), antibody VRC01 (n=6), or left them untreated (n=6) (Fig. 5A). While untreated mice showed stable viremia over the course of 4 weeks, mice treated with BG24 or VRC01 showed a comparable peak drop in average viral load of 0.54 and 0.57 log_10_ copies/ml, respectively (Fig. 5A). In both treatment groups, rebound of viremia occurred by 3 weeks post treatment initiation. To study viral escape mutations from BG24 *in vivo*, we performed single genome sequencing of HIV-1 envelope from mouse plasma of three BG24-treated mice at four weeks after therapy initiation (Fig. 5B). All 22 sequences obtained harbored one or more recurrent mutations in the D-loop, CD4-binding loop or V5-loop region, including well-known mutations N279K, N280D and G459D which have been associated with CD4-binding site antibody escape (*18, 34, 66*). 18 post-rebound sequences were obtained from three mice treated with VRC01 which showed a similar escape mutation profile with recurrent mutations also including N279K, N280D and G459D (Fig. 5B). Neutralization testing of BG24 and VRC01 on an extended HIV_YU2_ site mutant panel further confirmed their similar mutational sensitivity profile (Table S6). We conclude that BG24 has similar therapeutic efficacy to VRC01 in HIV_YU2_-infected humanized mice, highlighting that BG24-like antibodies retain activity *in vivo*.

**Fig. 5.**
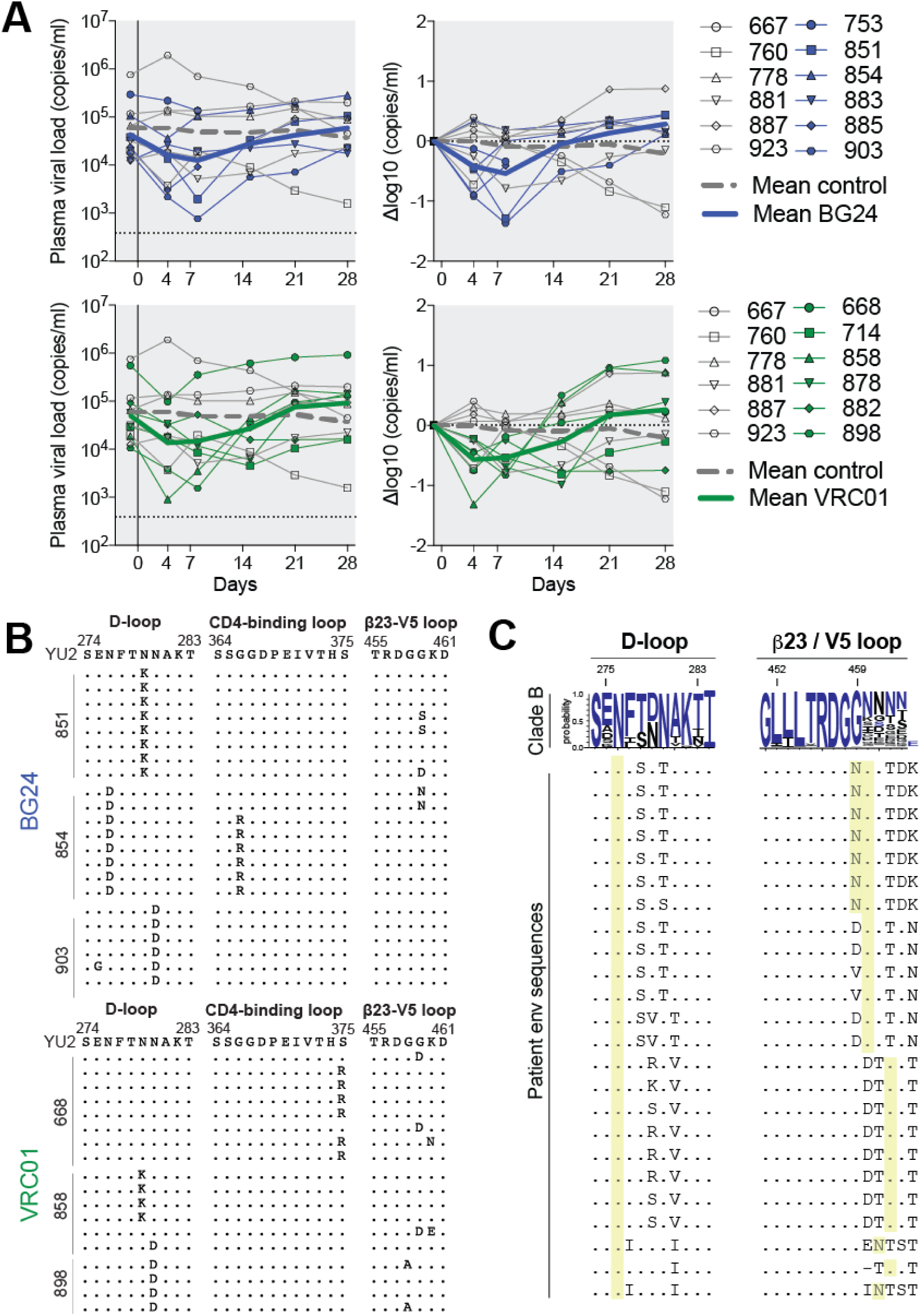
BG24 has comparable *in vivo* efficacy to VRC01 in HIV_YU2_-infected humanized mice. A) Antibody monotherapy of humanized mice infected with HIV_YU2_. Left graphs show absolute viremia (y-axis) in mice treated with antibody monotherapy (BG24, n=6, dark blue; VRC01, n=6, dark green) or untreated control mice (n = 6, grey) over the course of the experiment (x axis, days). Right graphs show relative log drop after initiation of antibody therapy (Δlog10 copies/mL). Thick blue/green and thick dashed gray lines indicate the mean viral load of treated and untreated mice, respectively. Mice were infected 3 weeks prior to therapy initiation and received 1 mg of IgG as a loading dose followed by twice-weekly administration of 0.5 mg for 3 weeks. The dotted line at the bottom indicates the limit of accuracy of the qPCR assay (384 copies/mL). Data from one experiment. B) Plasma HIV-1 Env sequences obtained 4 weeks after initiation of therapy from mice treated with BG24 (top) and VRC01 (bottom), respectively. Letters show amino acid mutations relative to the HIV_YU2_ molecular clone. Residues numbered according to HIV-1_HXB2_. C) HIV-1 Env sequences obtained from donor plasma RNA. Letters indicate amino\acid mutations compared with consensus clade B (blue letters) shown on top. Black letters in the consensus sequence indicate amino acids also observed at each position with lower frequencies. Yellow columns indicate potential N-linked glycosylation sites (PNGSs). Residues are numbered according to HIV-1_HXB2_.

### Single genome sequencing of 391370’s plasma Env

In order to better understand the viral context in which BG24 arose, and to investigate selective pressure exerted by BG24 on Env, we sequenced contemporaneous plasma envelope of subject 391370 using single genome sequencing. We obtained 24 intact full-length Env sequences, which all mapped conclusively as Clade B. As has been described in other elite neutralizers such as CH505 (*5, 67*), a high-level of diversity was evident in the Env quasispecies of 391370. The Env phylogeny segregated into two major branches that exhibited an average sequence difference of more than 20% of Env nucleotides (Fig. S8). Consistent with selective pressure being exerted by the BG24 family, we observed positive selection for known CD4-binding site escape mutations in the D-loop (D279K, D279R, A281T), and the b23-V5 loop region (G459D, addition of a glycan in V5), and all sequences carried a PNGS at 276_gp120_ (Fig. 5C).

### The role of somatic hypermutation in enhancing potency and breadth

Analysis of the BG24 paratope revealed 50% of the paratope surface to involve V-gene regions of both heavy and light chains, and an additional 22% attributed to CDR3 regions (Fig. 6A,B). Thus, only 28% of paratope residues were altered by somatic hypermutation, with the majority of somatic hypermutations occurring in CDR loops. Somatic mutations in FWR1 and CDRH1 mediated interprotomer gp120 contacts, while mutations in CDRL1 (including a 6 amino acid deletion) were necessary for accommodating the N276_gp120_-glycan at the CD4bs, consistent with other VRC01-class bNAbs (Fig. 6C-E). Interestingly, BG24 somatic hypermutation primarily modified CDRH2 and neighboring FWR sequence motifs to provide potential H-bonding to the CD4bs loop (Fig. 6F).

**Fig. 6.**
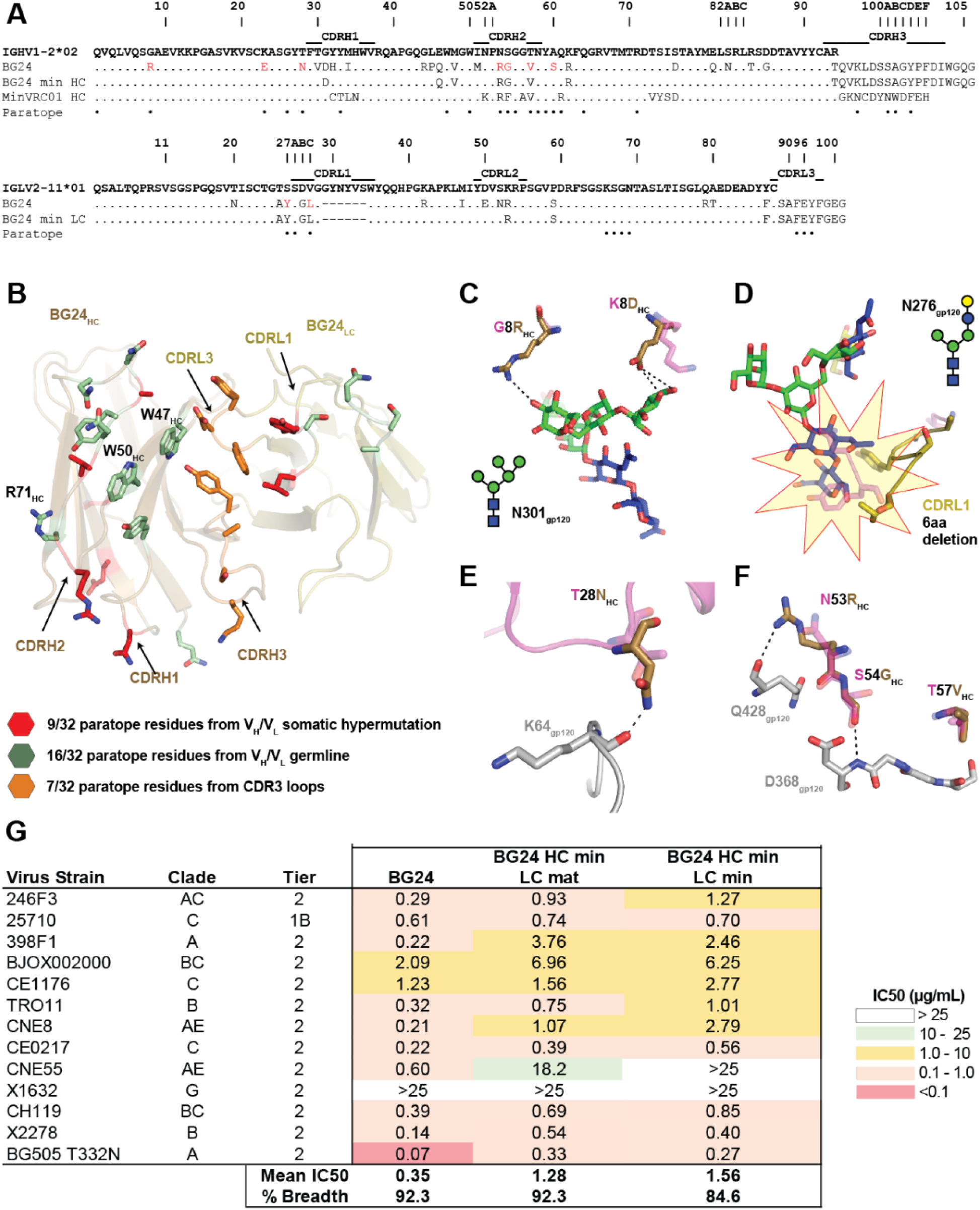
The role of somatic hypermutation in BG24 recognition of gp120. A) Sequence alignment of BG24 and BG24 minimally-mutated constructs with germline sequences. Somatic mutations compared to germline are shown with BG24 paratope residues derived from SHM shown as red. Complete paratope is labeled below sequence alignment. B) Paratope residues from germline V genes (green), somatic mutation (red) and CDR3 loops (orange) are shown as sticks on BG24. C-F) Representative mutations that increase contacts of mature BG24 with CD4bs. Model of germline BG24 (pink) was superposed with the BG24-BG505-101074 structure. F) Neutralization data of engineered minimally-mutated BG24 constructs against the global 12-strain viral panel. The average mean IC_50_s in µg/mL are shown from duplicate neutralization measurements.

To better define somatic hypermutations in BG24, we designed several heavy and light chain constructs to determine regions that played an essential role in its neutralization breadth and potency. Inclusion of the 6-residue CDRL1 deletion in the BG24 LC did not result in any appreciable binding or neutralization to Env isolates containing the N276_gp120_-glycan with the exception of the clade D isolate, 6405. Thus, all minimal constructs were constructed to maintain the N276_gp120_-glycan accommodating CDRL1 deletion (Fig. 6A), similar to the previously described MinVRC01 (*37*). Reversion of somatic hypermutations in the FWRs, CDRH1, and CDRL2, yielded a minimally-mutated BG24 construct that showed ∼85% neutralization breadth on the global 12-strain viral panel with a geometric mean IC_50_ of 1.56 µg/mL (Fig. 6A,G). This observation is consistent with longitudinal analysis of PCIN63 maturation where early convergence of CDRH2 motifs was critical to heterologous activity (*38*).

## Discussion

VRC01-class bNAbs have long been a target for rational vaccine design given their near pan-neutralizing activity across HIV-1 viral clades (*32, 68*). However, despite success in the isolation of numerous VRC01-class bNAbs, the prospect of eliciting such antibodies in vaccination efforts has been hampered by the high mutational count of these antibodies (*32, 61, 68*). Recent B-cell repertoire data suggests that the frequency of SHM observed in archetypal VRC01-class bNAbs ranks far above the average SHM frequency in the B-cell repertoires of HIV-naïve individuals (*69*). Indeed, it was suggested that such heavily mutated bNAbs might only arise in the context of HIV-1 infection and associated changes in normal B-cell selection processes. Easier blueprints for potent and broad CD4bs antibodies that might be more readily elicited by vaccination, therefore, remain an area of strong interest. Here, we studied the antibody response of a viremic controller and identified VRC01-class antibody BG24 that might represent one such easier blueprint for CD4-binding site based HIV vaccine design.

Our results indicate that there are exceptions to the commonly-held rule that high levels of somatic hypermutation are a pre-requisite for the breadth and potency of VRC01-class antibodies (*70, 71*). While it had been previously demonstrated that minimally mutated VRC01-class antibodies can be constructed through *in-silico* based design (*37*), only one naturally arising VRC01-class antibody lineage, the PCIN63 family, with a shorter maturation path and comparable neutralization activity to VRC01 has been described to date (*38*). Similarly, to the PCIN63 lineage, antibody BG24 shows low rates of SHM in a range of <15% on nucleotide level in both heavy and light chain, a range of mutation that might be more readily achievable by vaccination. Indeed, we found that only 28% of the BG24 paratope was modified by SHM and found key mutations to be focused within specific regions, in particular CDRH2 and heavy chain framework residues. In line with these findings and previous data (*56*), we were able to show that it is possible to construct even less mutated versions of BG24-type antibodies that exhibit high breadth and modest neutralization with <10% SHM on the amino acid level.

Taken together, BG24 and the PCIN63 lineage indicate that multiple shorter mutational pathways to VRC01-class type recognition of the CD4-binding site exist. Indeed, the two bNAbs arose in the context of different infecting viral clades, as PCIN63 arose in a Clade C infected donor, while BG24 arose in a Clade B infected donor. Moreover, while PCIN63 utilizes an IGK1-5*03 light chain and accommodates the N276_gp120_-glycan through CRDL1 flexibility, BG24 employs an IGLV2-11*01 light chain and accommodates the N276_gp120_-glycan through a 6 amino-acid CRDL1 deletion. In contrast to some other VRC01-class antibodies, neither of the two antibodies carries insertions or deletions in the heavy chain. Overall, the heavy chain sequences of these two antibodies are quite divergent with 29.2% difference on amino acid level in their V-gene portions despite both appearing to be derived from IGHV1-2*02 germline genes, suggesting that the trajectory of VRC01-class bNAb affinity maturation is not limited and can sample diverse sequences at key contact residues.

Providing support for the notion that such VRC01-class antibodies with short maturational pathways retain stability and potential for anti-HIV activity *in vivo,* we were able to demonstrate that BG24 had comparable therapeutic *in vivo* efficacy to VRC01 in humanized mice. We also did not find any indication of auto- or polyreactivity of BG24 suggesting that such phenomena might not be an impediment to BG24-type antibody induction. Moreover, engineered constructs that encoded neutralizing enhancing mutations in CDRH2, including the G54WHC mutation, also showed no polyreactivity, suggesting that improvement of such antibodies for therapeutic use can be achieved.

Structural studies of BG24 bound to a BG505 and DU422 Env trimers demonstrated that BG24 exhibited a conserved binding orientation relative to more mutated VRC01-class bNAbs, maintaining critical germline and CDR3 interactions, but established interprotomer contacts with the adjacent gp120. While the overall binding orientation of BG24 to Env was conserved, BG24 deviates from other VRC01-class antibodies in signature sequence features, including in particular the use of a Tyr instead of a Trp in the -5 position of the CDRH3 and also does not exhibit the consensus QQYEF CRDL3 motif of the VRC01-class antibodies that are derived from kappa germline genes. Indeed, BG24 belongs to the less frequently described group of VRC01-class antibodies that use a lambda light chain and is the second IGLV2-11*01 CD4bs bNAb lineage isolated to date (*51, 57*).

A recent study that assessed binding of B-cell receptors in the immune repertoire of HIV-naïve individuals to the CD4bs immunogen eOD-GT8 identified lambda light chain-using VRC01-class antibody precursors with a Tyr residue at the -5 position of CDRH3 (*54*), supporting the notion that BG24 precursors exist widely. In contrast to the previously dominating assumption that a Trp in CDRH3 position -5 is key, BG24’s properties actually demonstrate that a Tyr at the -5 position can even be favorable in specific instances, as we found that replacement of Tyr with a Trp reduced neutralization activity. This suggests that these naïve B-cells isolated from HIV-uninfected donors using eOD-GT8 or other VRC01-class germline-targeting immunogens may represent bona-fide VRC01-class bNAb precursors with potential to develop into BG24-type CD4-binding site bNAbs.

Collectively, our data suggests that BG24-type antibodies represent a potential target for CD4- binding site directed HIV vaccine design. Future studies will be required to explore whether boosting immunogens will be able to shepherd presumed BG24 B-cell precursors to reach broad and potent neutralization, in particular given that the N276_gp120_-glycan barrier still presents a major hurdle towards these efforts. Overall, the discovery of donor-derived minimally-mutated VRC01-class bNAbs and their intermediates raises the possibility that immunization schemes that might drive such responses are within reach.

## Materials and Methods

### Patient samples

Subject 391370 was a participant in The Ragon Institute of MGH, MIT and Harvard protocol ‘Host Genetics, Immunology and Virology of HIV’ from October 2005 to April 2008. The protocol was approved by the MGH IRB (2003P001678/MGH). Biological samples were obtained and analyzed under protocol MNU-0625 approved by the Rockefeller IRB. The subject is an African American male originally diagnosed with HIV-1 in 1990 who was always ART-naïve until the end of the study follow-up. B-cell sorting was done on two aliquots of a peripheral blood mononuclear cell (PBMC) sample from 2007. During the period of observation, viral loads ranged from 215 to 27,400 copies/ml (geometric mean: 795 copies/ml) and CD4^+^ T-cell counts from 352-495 cells/µl (mean: 413 cells/µl). Polyclonal IgG from subject 391370 was isolated from heat-inactivated plasma using Protein G Sepharose 4 Fast Flow (GE Healthcare). Purified IgG was buffer exchanged into Dulbecco’s Phosphate Buffered Saline (DPBS) and sterile-filtered prior to neutralization testing.

### Neutralization testing by TZM.bl and neutralization fingerprinting

A luciferase-based TZM.bl assay was used to measure the neutralizing activity of polyclonal IgG and monoclonal antibodies according to standard protocols (*72*). Each assay was performed at least in duplicates. To determine 50% (IC_50_) or 80% (IC_80_) inhibitory concentrations, 5-parameter curve fitting was utilized. Non-specific activity was detected by testing against murine leukemia virus (MuLV). Analysis of neutralization and graphing was done using the Antibody Database (v 2.0) (*73*). In order to determine the neutralization fingerprint of the polyclonal IgG/monoclonal antibodies, a panel of 20 HIV-1 strains (f61 panel) was used as described (*42, 74*).

### Single B-cell bait-sorting

Single B-cell sorting of IgG^+^ memory B-cells was done using BG505.SOSIP.664 as bait in a manner previously described with slight modifications (*44, 75, 76*). Avi-tagged BG505.SOSOP.664 (BG505.SOSIP.664.Avi) was produced in CHO cells and purified using a PGT145 affinity column as described (*76, 77*). BG505.SOSIP.664.Avi was biotinylated using the BirA-Ligase (Avidity) following the manufacturer’s instructions, and an aliquot of 5 µg of biotinylated BG505.SOSIP.664 was then freshly coupled to Streptavidin-PE in a volume of 10 µl DPBS immediately before sorting. Two independent sorts on 10 million PBMCs each were carried out. In the first sort, fluorescent staining of total PBMCs was done using CD3-PerCP-Cy5.5, CD14 PerCP-Cy5.5, CD335 PerCP-Cy5.5, CD606 PerCP-Cy5.5, CD19 BV421, CD20 BV421, IgG BV510, IgM BV605 and fluorescently-labelled BG505 bait. The staining for the second sort included the same antibodies with the addition of LIVE/DEAD Fixable Aqua. Stainings were performed for 30 mins at 4 °C, and cell sorting was done on a FACS Aria II. BG505-binding IgG^+^ memory B-cells were sorted directly into lysis buffer in 96-well plates (*78, 79*). Amplification of B-cell heavy and light chain variable regions was done as described (*41, 44*), and bands from positive wells were subjected to Sanger Sequencing. Analysis of obtained antibody gene sequences was done using IgBLAST and the international ImMunoGeneTics information system (IMGT). For recombinant production, antibody variable regions were cloned into human Igγ1-, Igκ or Igλ-expression vectors by sequence and ligation independent cloning (SLIC). Recombinant expression of antibodies was done using transient transfection of 293-6E cells followed by Protein G purification. Antibodies for neutralization, and *in vivo* studies were buffer exchanged into DPBS using Amicon Ultra centrifugal filters.

### Phylogenetic analysis of BG24 clonal members

The human IgV_H_1-2*02 allele sequence was sourced from the international ImMunoGeneTics information system (IMGT) (*80*). Heavy chain nucleotide sequences of the BG24 clonal family were aligned with the IgV_H_1-2*02 germline sequence in Geneious R8 (v8.1.9) using MUSCLE. The maximum-likelihood tree was then generated with the RAxML plugin (v 7.2.8) using a GTR Gamma model and the ‘Rapid Bootstrapping and search for best-scoring ML tree’ function with 100 bootstrap replicates. Formatting of the best-scoring ML tree was done using FigTree (v1.4.3).

### Autoreactivity and polyreactivity assays

Autoreactivity of antibody BG24 and the two reference antibodies 4E10 and 2F5 was evaluated with the commercially-available HEp-2 based assay NOVA Lite kit (Inova Diagnostics). Testing was performed at an IgG concentration of 25 µg/ml. Photographing of slides was done on a Leica DMI 6000 B with 800 ms exposure, Gain of 10 and an intensity of 100%. Measurements were performed in duplicate.

Baculovirus-based polyreactivity assays were conducted using ELISA detection of non-specific binding as described (*63*). Briefly, a solution of 1% baculovirus particles in 100mM sodium bicarbonate buffer pH 9.6 was absorbed onto the wells of a 384-well ELISA plate (Nunc Maxisorp) using a Tecan Freedom Evo liquid handling robot. The plate was incubated overnight at 4°C followed by a 1 h block at room temperature with PBS + 0.5% BSA. Purified IgGs (diluted to 1 µg/mL in PBS + 0.5% BSA) were added to the blocked assay plate and incubated for 3 hours at room temperature. Bound IgG was detected as the luminescence signal at 425 nm using an HRP-conjugated anti-human IgG (H&L) secondary antibody (Genscript) and SuperSignal ELISA Femto Maximum Sensitivity Substrate (Thermo Fisher Scientific).

### *In vivo* experiments

Mouse experiments were approved by the State Agency for Nature, Environment and Consumer Protection (LANUV) of North-Rhine Westphalia. NOD-Rag1^null^ IL2rg^null^ (NRG) mice were purchased from The Jackson Laboratory. NRG mice were bred and maintained at the Dezentrales Tierhaltungsnetzwerk Weyertal at University of Cologne. Assessment of the pharmacokinetics of BG24 was done in non-humanized NRG mice. Mice were injected intravenously with 250 µg of BG24 (n=6) via the tail vein. Facial vein bleedings were done on days 1,3,6,9 and 14 post injection, and serum levels of the antibodies were measured using a total IgG ELISA as previously described (*18*). To generate humanized mice for HIV-1 treatment experiments, an established protocol was followed with slight modifications (*18, 81*). In brief, sub-lethally irradiated 1-5 day old NRG mice were injected intra-hepatically with CD34^+^ hematopoietic stem cells (HSCs) and screened for humanization by flow cytometry 12-weeks post HSC injection. HSCs were purified from cord blood or perfused human placental tissues using magnetic bead based purification (Miltenyi). The stem cell isolation protocol was approved by the ethics committee of the Medical Faculty of the University of Cologne. All stem cell donors provided written informed consent.

For antibody treatment experiments, HIV-1 infection of humanized mice was performed intraperitoneally using HIV-1_YU2_ (*82, 83*). For viral load measurements, mice were bled from the facial vein into EDTA tubes (Sarstedt). Viral RNA was subsequently isolated from mouse plasma using the MinElute Virus Kit (Qiagen) on the QiaCube. Measurements of HIV-1 levels in plasma were done using an in-house quantitative PCR assay that amplifies a part of pol (*82*) using the TaqMan® RNA-to-Ct™ 1-Step Kit on a Roche LightCycler 480 Instrument II. The primers used in the qPCR were: 5’-TAATGGCAGCAATTTCACCA-3’ and 5’-GAATGCCAAATTCCTGCTTGA-3’, the probe was 5’-/56-FAM/CCCACCAAC/ZEN/ARGCRGCCTTAACTG/3IABkFQ/-3′. The assay was determined to have a limit of accuracy of 384 copies/ml (based on the standard curve used). Before starting treatment, viral loads of mice were measured two times. Only mice with viral loads of more than 4,000 copies/ml prior to treatment were used in experiments. Antibody injections were done subcutaneously. Treatment was initiated with a loading dose of 1 mg of each antibody, and mice subsequently received 0.5 mg of each antibody every 3 days for a total of 3 weeks.

### Single genome sequencing of mouse plasma HIV-1 env genes

Single genome sequencing of mouse plasma env genes was carried out as described previously (*44*). In brief, complementary DNA (cDNA) was generated from extracted mouse plasma RNA using primer YB383 5’-TTTTTTTTTTTTTTTTTTTTTTTTRAAGCAC-3’ and enzyme Superscript III (Invitrogen) according to manufacturer’s instructions. Synthesized cDNA was subsequently serially diluted and subjected to two rounds of nested PCR using Platinum Taq Green Hot Start (Thermo Fisher Scientific) and primers specifically adapted for amplification of envel of HIV_YU2NL4-3_ (1^st^ round primers: YB383 and YB50 5’-GGCTTAGGCATCTCCTATGGCAGGAAGAA-3’; 2^nd^ round primers YB49 5’-TAGAAAGAGCAGAAGACAGTGGCAATGA-3’, YB52 5’-GGTGTGTAGTTCTGCCAATCAGGGAAGWAGCCTTGTG-3’). Bands of proper size from amplifications with less than 30% efficiency were PCR-purified using the Nucleospin Gel and PCR-Clean Up kit (Macherey Nagel, 740609.250) and then Sanger sequenced. Assembly of Env sequences was done using the Geneious 8.1.9 (Biomatters) de-novo assembly tool. Sequences with full coverage of gp160 Env were used in downstream analyses.

### Single genome sequencing of patient HIV-1 Env genes

Single HIV-1 genomes encoding HIV-1 gp160 were amplified from patient plasma according to a previously published protocol (*84–86*). In brief, viral RNA was isolated from patient plasma using the Virus Mini Spin Kit on a QiaCube. Isolated viral RNA was then used to generate complementary DNA (cDNA) using Superscript III according to manufacturer’s instructions with the primer envB3out (5’– TTGCTACTTGTGATTGCTCCATGT-3’). The HIV-1 env gene was then amplified through a nested PCR approach using Platinum Taq. 1^st^ round primers were: envB5out 5’-TAGAGCCCTGGAAGCATCCAGGAAG-3’ and envB3out 5’-TTGCTACTTGTGATTGCTCCATGT-3’; 2^nd^ round primers were: envB5in 5’ – TTAGGCATCTCCTATGGCAGGAAGAAG-3’ and envB3in 5’–GTCTCGAGATACTGCTCCCACCC-3’. PCRs were carried out using serial dilutions of cDNA to obtain a range in which less than 30% of wells generated a band. Positive wells from amplifications that yielded less than 30% of bands were subjected to library preparation with the Nextera DNA Amplification Kit. Env libraries were sequenced on an Illumina MiSeq (2x 150 bp Nano Kit) and assembled to the best HIV-1 Env reference sequence from HIV Blast using an in-house pipeline (*84*). Only intact Env sequences with a maximum of one ambiguity were used in downstream analyses. To generate the maximum-likelihood tree of subject 391370’s plasma env sequences, env nucleotide sequences were aligned in Geneious R8 (v8.1.9) using ClustalW. The maximum-likelihood tree was then generated with the RAxML plugin (v 7.2.8) using a GTR Gamma model and the ‘Rapid Bootstrapping and search for best-scoring ML tree’ function with 100 bootstrap replicates. The best-scoring ML tree was formatted using FigTree (v1.4.3).

### Protein expression and purification for structural studies

Fabs and IgGs used in this study were produced as described (*44*). Briefly, Fabs and IgGs were expressed by transiently transfecting Expi293 cells with vectors encoding the appropriate heavy and light chain genes. Secreted Fabs or IgGs were purified from cell supernatants using Ni^2+^-NTA (Fabs) or Protein A affinity chromatography (IgGs) followed by size exclusion chromatography (SEC) with a Superdex200 16/60 column (Cytiva). Purified proteins were concentrated and maintained at 4 °C in storage buffer (20 mM Tris pH 8.0, 150 mM NaCl, 0.02% sodium azide).

Genes encoding soluble BG505 SOSIP.664 or DU422 SOSIP.664 gp140 trimers were stably expressed in Chinese hamster ovary cells as described (*77, 87*). Secreted Env trimers expressed in the absence of glycosylation inhibitors were isolated from cell supernatants using PGT145 immunoaffinity chromatography by covalently coupling PGT145 IgG monomer to an activated-NHS Sepharaose column (Cytiva) as described (*88*). Trimers were eluted using 3M MgCl_2_, dialyzed into storage buffer, and purified using a Superdex200 16/60 column (Cytiva) against the same buffer. Peak fractions pertaining to SOSIP trimers were pooled and repurified using the same column and buffer conditions. Individual fractions were stored separately at 4 °C.

### Crystal structures of BG24_S60A_ Fab and a BG24_S60A_-BG505-10-1074 complex

A complex of BG24_S60A_-BG505-101074 was assembled by incubating purified BG24_S60A_ Fab with BG505 SOSIP.664 trimer at a 3:1 Fab:gp120-protomer molar ratio. Following overnight incubation at RT, 10-1074 Fab was incubated with the complex at a 3:1 Fab:gp120-protomer molar ratio for 5 h, and complexes were purified from unbound Fab by SEC on a Superose 6 10/300 column (Cytiva) run in 20 mM Tris pH 8 and 100 mM NaCl. Purified complexes were concentrated to 5-10 mg/mL by centrifugation with a 100 kDa concentrator (Millipore). Unliganded BG24_S60A_ Fab was concentrated to 10-15 mg/mL by centrifugation with a 30-kDa concentrator (Millipore).

Initial matrix crystallization trials against 576 conditions were performed at room temperature using the sitting drop vapor diffusion method by mixing equal volumes of protein sample and reservoir using a TTP LabTech Mosquito robot and commercially-available screens (Hampton Research and Qiagen). Initial hits were optimized and crystals for unliganded BG24_S60A_ were obtained in 0.25 M potassium chloride, 18% polyethylene glycol (PEG) 3350 at 20 °C. Crystals for the BG24_S60A_-BG505-101074 complex were obtained in 0.1 M Bis-Tris pH 6.5, 20% PEG1500. Crystals were cryo-protected stepwise to 20% glycerol before being cryopreserved in liquid nitrogen.

X-ray diffraction data were collected for both samples at the Stanford Synchroton Radiation Lightsource (SSRL) beamline 12-2 on a Pilatus 6M pixel detector (Dectris). Data from a single crystal were indexed and integrated in XDS (*89*) and merged with AIMLESS in the *CCP4* software suite (*90*). The unliganded BG24_S60A_ structure was determined by molecular replacement in PHASER (*91*) using a single search with coordinates of the VRC-PG20 Fab (PDB 4LSU) after removal of CDR loops. Coordinates from a refined BG24_S60A_ model were used in combination with a gp140-10-1074 complex (PDB 5T3Z) as search models for the BG24_S60A_-BG505-101074 complex data. Models generated by molecular replacement were refined using B-factor refinement in Phenix (*92*), followed by several cycles of manual building with B factor sharpening in Coot (*93*). For the 2.0Å BG24_S60A_ Fab structure, TLS refinement was also performed.

### Cryo-EM sample preparation

A complex of BG24_CDR2-v2_-DU422-101074 was assembled by incubating purified BG24_CDR2-v2_ Fab with DU422 SOSIP.664 trimer at a 1.2:1 Fab:gp120-protomer molar ratio. Following overnight incubation at RT, 10-1074 Fab was incubated with the complex at a 1.2:1 Fab:gp120-protomer molar ratio for 5 h. BG24_CDR2-v2_-DU422-101074 complexes were concentrated to 1-2 mg/ml in 20 mM Tris pH 8 and 100 mM NaCl, and 3 µl was added to Quantifoil R1.2/1.3 300 mesh copper grid (Electron Microscopy Services) that had been freshly glow-discharged using a PELCO easiGlow (Ted Pella). Samples were immediately vitrified in 100% liquid ethane using a Mark IV Virtoblot (ThermoFisher) by blotting for 3-4s with Whatman No. 1 filter paper at 20°C and 100% relative humidity.

### Cryo-EM data collection and processing

Single-particle cryo-EM data were collected on a Talos Arctica transmission electron microscope (ThermoFisher) operating at 200 kV, using a 3x3 beam image shift pattern with SerialEM automated data collection software (*94*). Movies were collected on a Gatan K3 Summit direct electron detector (DED) operating in counting mode at a nominal magnification of 45,000x (super-resolution 0.4345 Å/pixel) using a defocus range of -1.0 µm to -2.5 µm. Movies were collected with an 3.6 s exposure time with a rate of 13.5 e^-^/pix/s, which resulted in a total dose of ∼60 e-/Å^2^ over 40 frames.

Data processing was conducted as previously described (*44*). Briefly, movies were motion corrected and doseweighted using MotionCor2 in RELION-3 (*95*). Non-dose weighted summed images were used for CTF determination using Gctf (*96*), and reference-free particle picking was achieved using Laplacian-of-Gaussian filtering in RELION-3 (*95*). An initial stack of 455,671 particles were extracted from 1,180 dose-weighted micrographs and subjected to reference-free 2D classification. A total of 310,246 particles corresponding to class averages that displayed secondary-structural elements and represented views different views of Fab bound Env-trimer were extracted and re-centered prior to heterogenous ab inito mod el generation using cryoSPARC v2.2 (*97*).

The generated volume was low-passed filtered to 60 Å and used as an initial model for 3D auto-refinement in RELION-3 (C1 symmetry, k=8). After 25 iterations, a soft mask was generated from the highest-resolution model (5-pixel extension, 10-pixel soft cosine edge), and used in an additional round of 3D classification. This procedure yielded a particle stack of 248,600 particles that was homogenously refined in Relion using a soft mask in which Fab constant domains were masked out. 3D classification was repeated without alignments and 204,220 particles were subjected to particle polishing, CTF refinement, and subsequent rounds of homogenous refinement with C3 symmetry applied. Refinement procedures produced a final estimated global resolution of 3.5Å Å according to gold-standard FSC (*98*).

### Modeling and refinement of cryo-EM structures

For the final reconstruction of BG24_CDR2-v2_-DU422-10-1074, initial coordinates were generated by docking a refined BG24_S60A_-BG505-10-1074 reference model (this work) into the cryo-EM density using UCSF Chimera v1.13 (*99*). After sequence matching to DU422 gp140, initial models were refined into the EM maps using one round of rigid body, morphing, and simulated annealing followed by subsequent rounds of B-factor refinement in Phenix (*92*). Models were manually built following iterative rounds of real-space and B-factor refinement in Coot (*93*) and Phenix (*92*) with secondary structure restraints. Modeling of glycans was achieved by interpreting cryo-EM density at PNGS in Coot using a map with a -150 Å^2^ B-factor sharpening value, contoured at 3σ due to the lower resolution of glycans at the periphery of the structure. Validation of model coordinates was performed using MolProbity (*100*) and Privateer (*101*).

### Structural and bioinformatic analyses

Superpositions and figures were rendered using PyMOL (Version 1.5.0.4 Schrodinger, LLC), and protein electrostatic calculations were done using APBS and PDB2PQR webservers (*102*). Buried surface areas (BSAs) were determined with PDBePISA using a 1.4Å probe (*103*). Potential hydrogen bonds were assigned using a distance of <3.6Å and an A-D-H angle of >90°, while the maximum distance allowed for a van der Waals interaction was 4.0Å. Putative H-bonds, van der Waals assignments and total BSA should be considered tentative, owing to the relatively low structure resolutions. Computational analysis of neutralization panel data (Table S7) was done as previously described (*73*).

## Acknowledgments

We thank members of the Bjorkman, Nussenzweig and Klein labs for helpful discussions. B-cell sorting of donor 391370 was performed with help of Kristie Gordon and Gaëlle Breton. Neutralization fingerprinting analysis was performed by Ivelin Georgiev. Sequencing analysis of antibody sequences and patient plasma env genes was supported by Joy Pai. BG505 SOSIP.664-Avi, the CHO cell lines expressing the BG505 SOSIP.664 v4.1 trimer and DU422 SOSIP.664 v4.1 trimer, and the BG505.T332N gp160 expression plasmid were kind gifts of Albert Cupo, John P. Moore and Rogier W. Sanders. We thank Dr. Jost Vielmetter, Pauline Hoffman, and other members of the Protein Expression Center in the Beckman Institute at Caltech for expression assistance. Structural studies were assisted by the Caltech Molecular Observatory (Dr. Jens Kaiser, director) and the Biological and Cryogenic Transmission Electron Microscopy Center at Caltech (Drs. Andrey Malyutin and Songye Chen, directors). This work was supported by the National Institute of Allergy and Infectious Diseases of the National Institutes of Health Grant HIVRAD P01 AI100148 (to P.J.B. and M.C.N.), National Institutes of Health Grant NIH P50 AI150464 (to P.J.B.), the Bill and Melinda Gates Foundation Collaboration for AIDS Vaccine Discovery Grants OPP1124068 (to M.C.N. and P.J.B.) and 1146996 (M.S.S.), NIH Center for HIV/AIDS Vaccine Immunology and Immunogen Discovery (CHAVI-ID) 1UM1 AI100663-01 (M.C.N.), the European Research Council (ERC-StG639961) (F.K.), the German Center for Infection Research (DZIF) (F.K.). C.O.B was supported by the Hanna Gray Fellowship Program from the Howard Hughes Medical Institute and the Postdoctoral Enrichment Program from the Burroughs Wellcome Fund. T.S. was supported by the Ernst Jung Career Advancement Award for Medical Research, grant UL1 TR001866 from the National Center for Advancing Translational Sciences (NCATS, NIH Clinical and Translational Science Award (CTSA) program), and DZIF grant TI 07.002. We thank the Gordon and Betty Moore and Beckman Foundations for gifts to Caltech to support the Molecular Observatory and electron microscopy. M.C.N. and B.D.W. are HHMI investigators.

## Author Contributions

C.O.B., T.S., M.C.N., and P.J.B., conceived the study. T.S. and J.G. performed B-cell sorting and antibody cloning of BG24 family members. C.O.B., Y.E.L., N.S-T., and K.E.H-T, performed protein purification and structural studies. P.N.P.G., K.E.H-T., A.P.W., and C.O.B., performed *in vitro* 12-strain neutralization assays on engineered BG24 constructs and analyzed the data. C.O.B., and N.S-T., conducted structural studies. T.S. and H.G. performed antibody *in vivo* experiments. P.S. conducted neutralization assays on viral mutants and envelope sequencing from mouse plasma. J.C.C.L. and T.S. conducted env sequencing from patient plasma. M.S.S. carried out neutralization testing on the 126-virus cross-clade panel. J.F.S. and M.S.S. conducted plasma screening of HIV controller cohort. A.P.T. and B.D.W. collected and provided patient samples from the HIV controller cohort. C.O.B., T.S., M.C.N., and P.J.B., wrote the manuscript with contribution from all authors.

## Data and materials availability

Nucleotide sequences of BG24 antibody family members and plasma env sequences from subject 391370 have been deposited in GenBank (Accession Numbers XXX-XXX). The atomic models of the unliganded BG24_S60A_ Fab and BG24-BG505–10-1074 complex have been deposited in the Protein Data Bank (PDB; http://www.rcsb.org/) under accession codes PDB 7UCE and 7UCF, respectively. Coordinates for the atomic model and cryo-EM map generated from cryo-EM studies of the BG24_CDR2-v2_-DU422-101074 complex has been deposited at the PDB and the Electron Microscopy Databank (EMDB, http://www.emdataresource.org/) under codes PDB 7UCG and EMD-26443, respectively. All data are available in the main text or the supplementary materials.

