## Supplemental Data for "A naturally arising broad and potent CD4-binding site antibody with low somatic mutation"

**This PDF file includes:**

Figs. S1 to S8
Tables S1 to S6

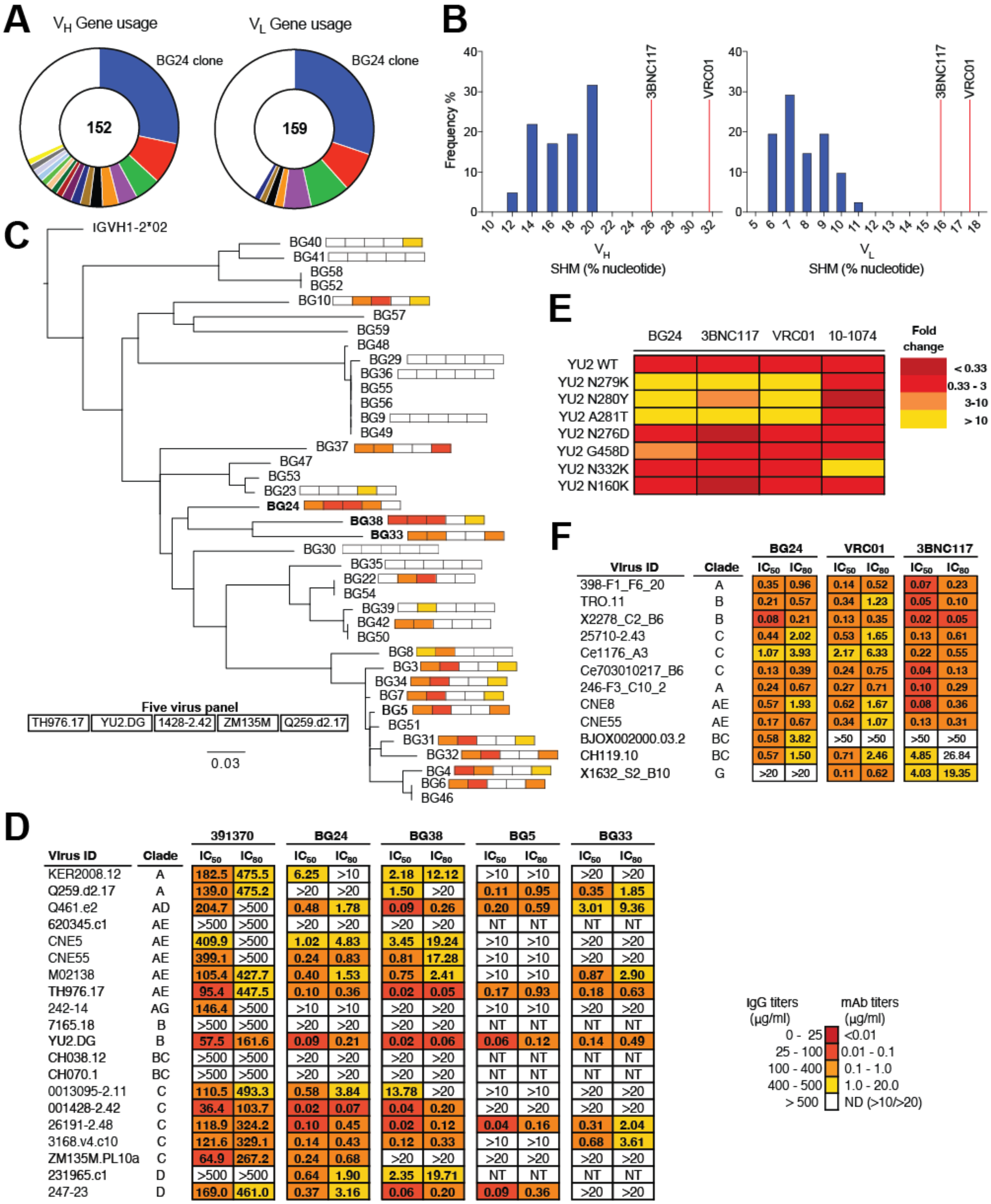

**D**

| Virus ID | Clade | 391370 IC50 | 391370 IC80 | BG24 IC50 | BG24 IC80 | BG38 IC50 | BG38 IC80 | BG5 IC50 | BG5 IC80 | BG33 IC50 | BG33 IC80 |
|---|---|---|---|---|---|---|---|---|---|---|---|
| KER2008.12 | A | 182.5 | 475.5 | 6.25 | >10 | 2.18 | 12.12 | >10 | >10 | >20 | >20 |
| Q259.d2.17 | A | 139.0 | 475.2 | >20 | >20 | 1.50 | >20 | 0.11 | 0.95 | 0.35 | 1.85 |
| Q461.e2 | AD | 204.7 | >500 | 0.48 | 1.78 | 0.09 | 0.26 | 0.20 | 0.59 | 3.01 | 9.36 |
| 620345.c1 | AE | >500 | >500 | >20 | >20 | >20 | >20 | NT | NT | NT | NT |
| CNE5 | AE | 409.9 | >500 | 1.02 | 4.83 | 3.45 | 19.24 | >10 | >10 | >20 | >20 |
| CNE55 | AE | 399.1 | >500 | 0.24 | 0.83 | 0.81 | 17.28 | >10 | >10 | >20 | >20 |
| M02138 | AE | 105.4 | 427.7 | 0.40 | 1.53 | 0.75 | 2.41 | >10 | >10 | 0.87 | 2.90 |
| TH976.17 | AE | 95.4 | 447.5 | 0.10 | 0.36 | 0.02 | 0.05 | 0.17 | 0.93 | 0.18 | 0.63 |
| 242-14 | AG | 146.4 | >500 | >10 | >10 | >20 | >20 | >10 | >10 | >20 | >20 |
| 7165.18 | B | >500 | >500 | >20 | >20 | >20 | >20 | NT | NT | NT | NT |
| YU2.DG | B | 57.5 | 161.6 | 0.09 | 0.21 | 0.02 | 0.06 | 0.06 | 0.12 | 0.14 | 0.49 |
| CH038.12 | BC | >500 | >500 | >20 | >20 | >20 | >20 | NT | NT | NT | NT |
| CH070.1 | BC | >500 | >500 | >20 | >20 | >20 | >20 | NT | NT | NT | NT |
| 0013095-2.11 | C | 110.5 | 493.3 | 0.58 | 3.84 | 13.78 | >20 | >10 | >10 | >20 | >20 |
| 001428-2.42 | C | 36.4 | 103.7 | 0.02 | 0.07 | 0.04 | 0.20 | >20 | >20 | >20 | >20 |
| 26191-2.48 | C | 118.9 | 324.2 | 0.10 | 0.45 | 0.02 | 0.12 | 0.04 | 0.16 | 0.31 | 2.04 |
| 3168.v4.c10 | C | 121.6 | 329.1 | 0.14 | 0.43 | 0.12 | 0.33 | >10 | >10 | 0.68 | 3.61 |
| ZM135M.PL10a | C | 64.9 | 267.2 | 0.24 | 0.68 | >20 | >20 | >20 | >20 | >20 | >20 |
| 231965.c1 | D | >500 | >500 | 0.64 | 1.90 | 2.35 | 19.71 | NT | NT | NT | NT |
| 247-23 | D | 169.0 | 461.0 | 0.37 | 3.16 | 0.06 | 0.20 | 0.09 | 0.36 | >20 | >20 |

**F**

| Virus ID | Clade | BG24 IC50 | BG24 IC80 | VRC01 IC50 | VRC01 IC80 | 3BNC117 IC50 | 3BNC117 IC80 |
|---|---|---|---|---|---|---|---|
| 398-F1_F6_20 | A | 0.35 | 0.96 | 0.14 | 0.52 | 0.07 | 0.23 |
| TRO.11 | B | 0.21 | 0.57 | 0.34 | 1.23 | 0.05 | 0.10 |
| X2278_C2_B6 | B | 0.08 | 0.21 | 0.13 | 0.35 | 0.02 | 0.05 |
| 25710-2.43 | C | 0.44 | 2.02 | 0.53 | 1.65 | 0.13 | 0.61 |
| Ce1176_A3 | C | 1.07 | 3.93 | 2.17 | 6.33 | 0.22 | 0.55 |
| Ce703010217_B6 | C | 0.13 | 0.39 | 0.24 | 0.75 | 0.04 | 0.13 |
| 246-F3_C10_2 | A | 0.24 | 0.67 | 0.27 | 0.71 | 0.10 | 0.29 |
| CNE8 | AE | 0.57 | 1.93 | 0.62 | 1.67 | 0.08 | 0.36 |
| CNE55 | AE | 0.17 | 0.67 | 0.34 | 1.07 | 0.13 | 0.31 |
| BJOX002000.03.2 | BC | 0.58 | 3.82 | >50 | >50 | >50 | >50 |
| CH119.10 | BC | 0.57 | 1.50 | 0.71 | 2.46 | 4.85 | 26.84 |
| X1632_S2_B10 | G | >20 | >20 | 0.11 | 0.62 | 4.03 | 19.35 |

**Fig. S1. Analysis of BG24 antibody family.**

A) Pie charts showing the number of heavy and light chain sequences recovered from $IgG^+$ memory B-cells, with individual clones represented by slices. HC/LC members of the $V_H$1-2*02/$V_L$2-11*01 using BG24 clone are shown as dark blue. White slices represent singlet sequences.

B) Frequency distribution of the % nucleotide somatic mutation from the germline variable gene in BG24 clonal family members (blue bars) for heavy chains (left) and light chains (right). Red lines denote the percentage nucleotide mutation of monoclonal antibodies 3BNC117 and VRC01.

C) Maximum-likelihood phylogenetic tree of nucleotide sequences of BG24 clone members. Colored boxes indicated mean $IC_{50}$ neutralization values against a 5-virus panel. Names of clone members that were selected for further testing are bolded.

D) Neutralization data of BG24, BG38, BG5, BG33 and subject 391370's serum IgG against virus members of the f61 virus panel. The mean $IC_{50}$s in µg/mL are shown from duplicate neutralization measurements.

E) BG24, 3BNC117, VRC01, and 10-1074 neutralizing activity against a panel of $HIV_{YU2}$ site mutant pseudoviruses. Displayed is the fold change in $IC_{50}$ relative to neutralization of $HIV_{YU2}$ wildtype. Data from duplicate neutralization measurements.

F) Neutralization data of BG24, 3BNC117, and VRC01 against the global 12-strain viruspanel. The mean $IC_{50}$s in µg/mL are shown from duplicate neutralization measurements.

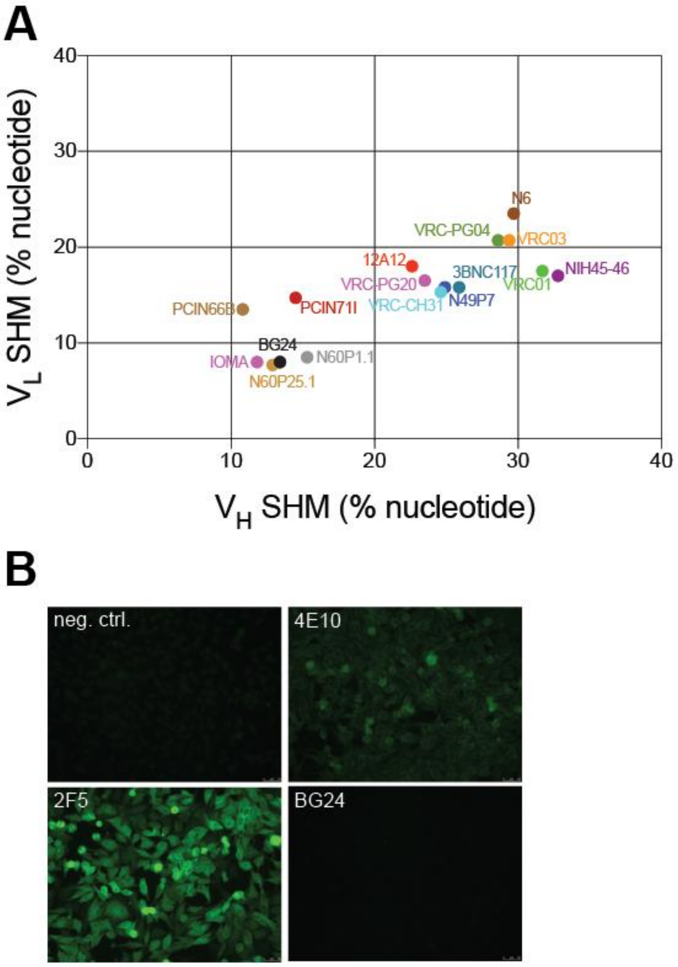

**Fig. S2. Mutational analysis of BG24 family members.**

A) Somatic mutation analysis of VRC01-like bNAbs, shown as % nucleotide changes relative to germline variable gene sequence.

B) HEp2 assay testing for potential autoreactivity of BG24 in comparison to reference antibodies 4E10 and 2F5. Testing carried out in duplicate.

**A**

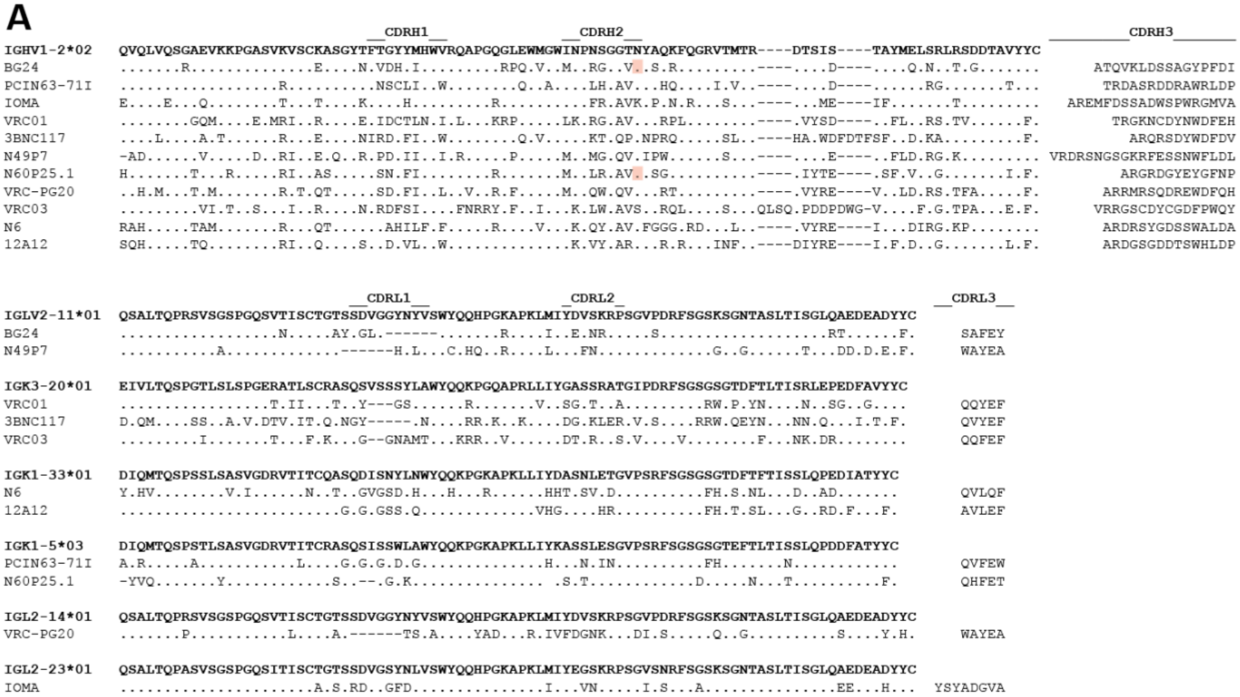

**B**

| Virus Strain | Clade | Tier | BG24 | BG24 S60A | BG24 S60T |
|---|---|---|---|---|---|
| 246F3 | AC | 2 | 0.29 | 0.26 | 1.28 |
| 25710 | C | 1B | 0.61 | 0.43 | 4.75 |
| 398F1 | A | 2 | 0.22 | 0.11 | 1.35 |
| BJOX002000 | BC | 2 | 2.10 | 1.50 | 24.1 |
| CE1176 | C | 2 | 1.23 | 0.99 | 10.6 |
| TRO11 | B | 2 | 0.32 | 0.22 | 1.62 |
| CNE8 | AE | 2 | 0.21 | 0.16 | 0.42 |
| CE0217 | C | 2 | 0.22 | 0.16 | 0.83 |
| CNE55 | AE | 2 | 0.60 | 0.37 | 3.69 |
| X1632 | G | 2 | >25 | >25 | >25 |
| CH119 | BC | 2 | 0.39 | 0.36 | 2.10 |
| X2278 | B | 2 | 0.14 | 0.09 | 0.85 |
| BG505 T332N | A | 2 | 0.07 | 0.08 | 0.60 |
| | | Mean IC50 | 0.35 | 0.26 | 2.04 |
| | | % Breadth | 92.3 | 92.3 | 92.3 |

IC50 (µg/mL): > 25 | 10 - 25 | 1.0 - 10 | 0.1 - 1.0 | <0.1

**Fig. S3. Sequence alignment of VRC01-class antibodies and N58$_{HC}$ glycan neutralization analysis of antibody BG24.**

A) Sequence alignment of VRC01-class bNAbs with their deduced germline V genes. CDR loops are indicated. Red boxes indicate potential N-linked glycosylation sites (PNGSs).

B) Neutralizing activity of BG24 and BG24 constructs ±N58$_{HC}$-glycan against the global 12-strain virus panel. Mean $IC_{50}$ values from duplicate measurements are shown.

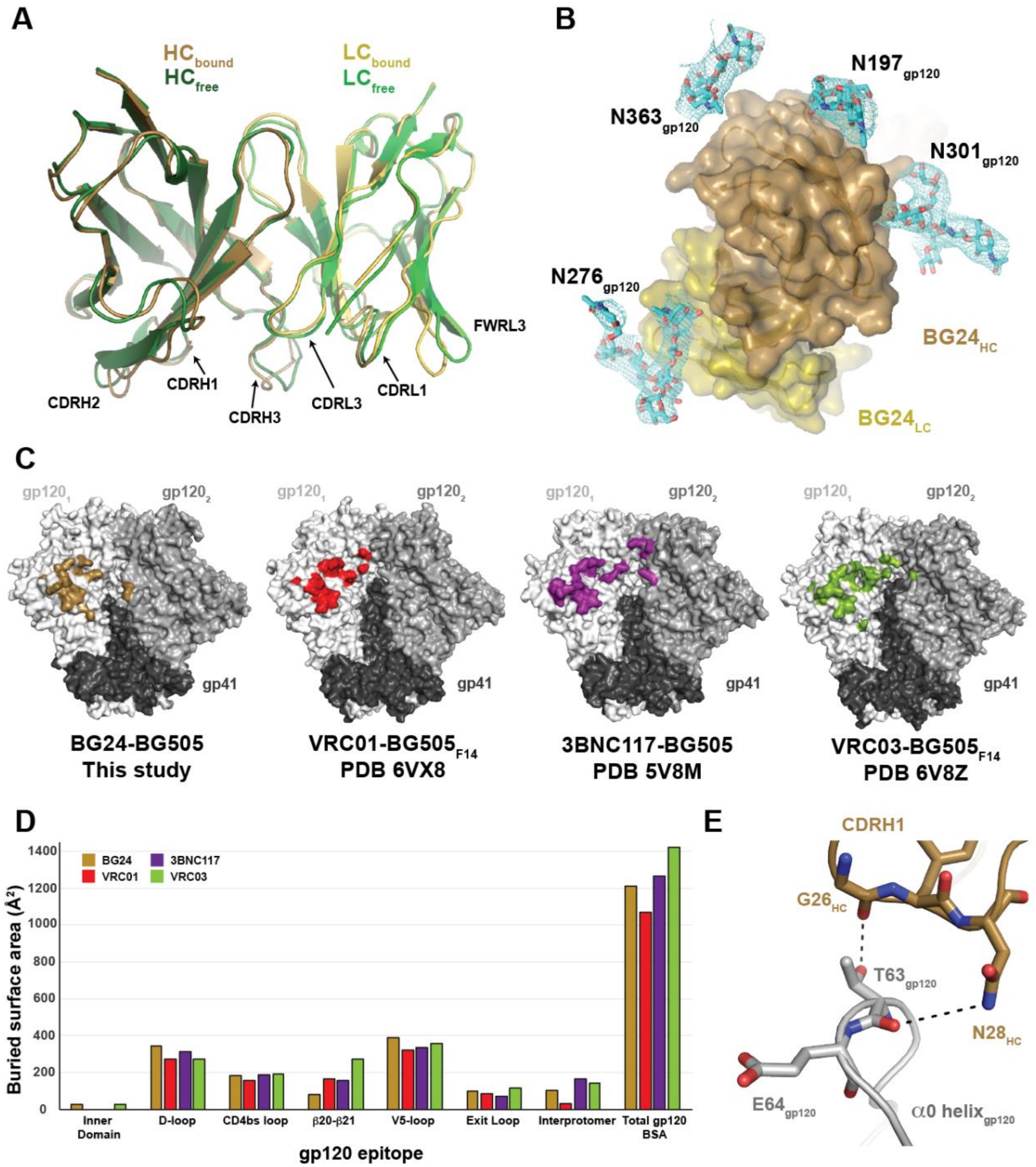

**Fig. S4. Characterization of BG24 epitope on BG505 gp120.**

A) Superposition of variable domains of unliganded (green) or BG505-bound (brown) BG24. CDR-loops and FWRs are indicated.

B) Surface representation of BG24 in the BG505-bound structure with observed gp120 CD4bs-glycans (cyan sticks). Glycan electron density (cyan mesh) contoured at 1.5σ.

C) Epitopes of CD4bs bNAbs on Env SOSIP.664 trimer structures indicated by colored surfaces on gp120 protomers. PDBs used for epitope calculations for each bNAb are shown.

D) Buried surface areas (BSA) of BG24 and VRC01-class bNAb contacts on the Env trimer surface, color-coded and derived from structures shown in panel C. BSA calculations for CD4bs regions are shown.

E) Stick representation of potential contacts between the CDRH1 of BG24 and α0 residues of the adjacent gp120 protomer of Env SOSIP.664. Black dashed lines indicate potential H-bond interactions.

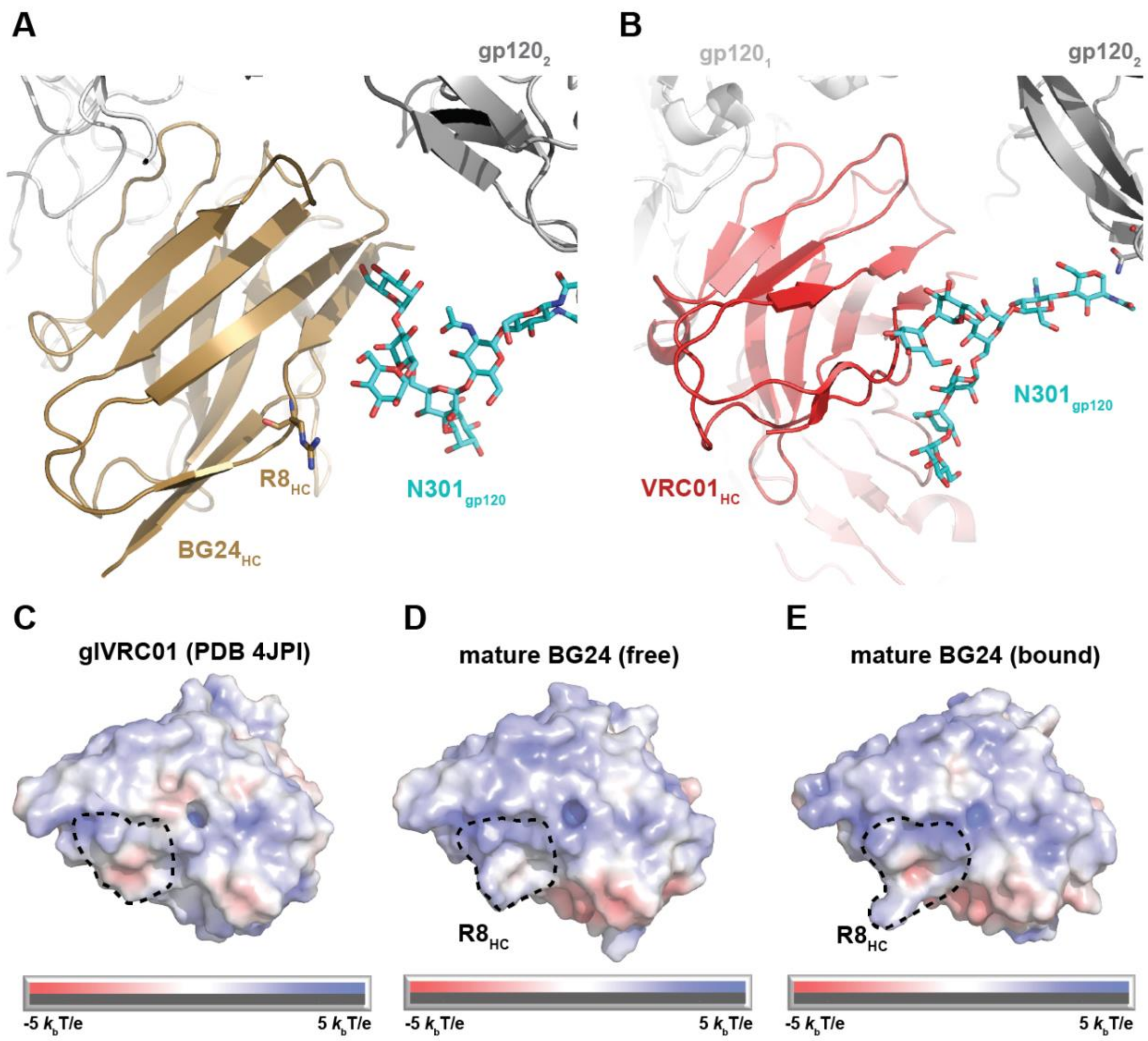

**Fig. S5. Contribution of $N301_{gp120}$-glycan to BG24 maturation.**

A,B) Cartoon and stick representation of potential contacts between the $N301_{gp120}$-glycan (cyan stick) from the neighboring gp120 protomer (gray) and A) BG24 (brown), or B) VRC01 (red, PDB 6VX8).

C-E) Surface representation of C) glVRC01, D) unliganded BG24, and E) BG505-bound BG24 colored by electrostatic surface potentials with red indicating negative electrostatic potential and blue representing positive electrostatic potential. Dotted line indicates potential $N301_{gp120}$-glycan footprint on antibody.

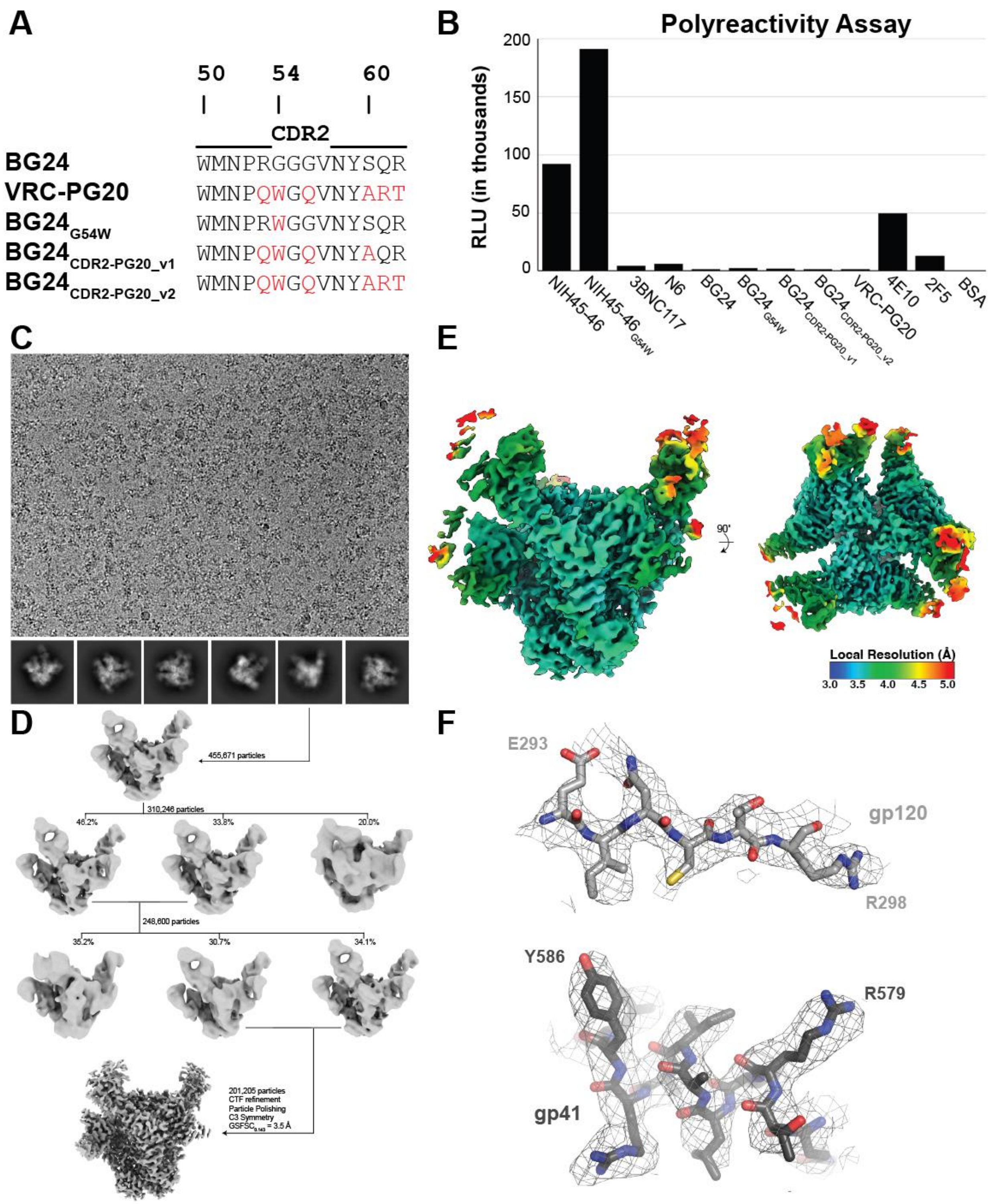

**Fig. S6. Analysis of CDRH2 loop-modified BG24 variants.**

A) Sequence alignment of CDRH2 amino acid residues of BG24, VRC-PG20, and engineered Phe43 pocket-filling BG24 constructs.

B) Polyreactivity ELISA-based assay detecting non-specific binding of a panel of HIV-1 bNAbs and a control protein (BSA) to a baculovirus extract.

C) Representative micrograph and 2D-class averages of $BG24_{CDR2\text{-}PG20\text{-}v2}$ – DU422 – 10-1074 complex.

D) 3D classification and refinement strategy for the $BG24_{CDR2\text{-}PG20\text{-}v2}$ – DU422 – 10-1074 complex.

E) Local resolution estimations calculated using ResMap in Relion.

F) Representative electron density contoured at $7\sigma$ for gp41 and gp120 residues (sticks).

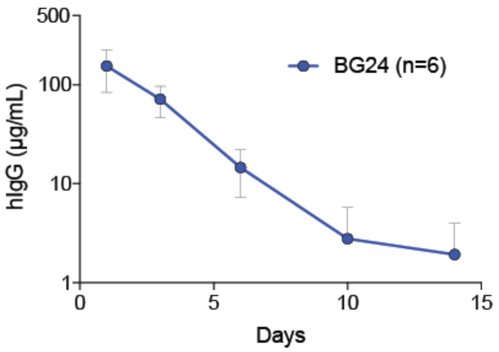

**Figure S7. Pharmacokinetic analysis of BG24 in NRG mice.** Analysis of BG24 concentration over time in serum of NRG mice after intravenous injection of 250 µg BG24 IgG1. Shown is the mean concentration with standard deviation of six mice from 2 independent experiments of 3 mice each.

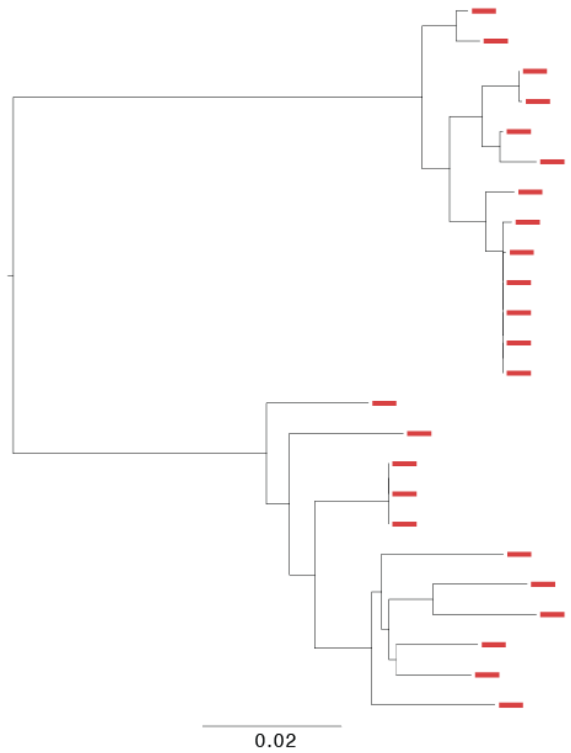

**Figure S8. Phylogenetic analysis of HIV-1 env from subject 391370's plasma.** Maximum-likelihood phylogenetic tree of subject 391370's plasma env nucleotide sequences. Tree is midpoint rooted.

**Table S1. Plasma neutralizing activity of subject 391370 and other HIV-1 infected subjects from the HIV Controller Cohort against an early cross-clade pseudovirus panel.**

| Virus | Clade/Tier / Subject | Patient 3 | 391370 | Patient 10 | Patient 8 | Patient 7 | Patient 9 |
|---|---|---|---|---|---|---|---|
| | Breadth | 100% | 93% | 93% | 93% | 93% | 87% |
| | Geomean ID80 | 74 | 132 | 102 | 97 | 82 | 186 |
| SF162.LS | B-Tier 1A | 429 | 2381 | 2381 | 2539 | 4417 | 4973 |
| BaL.26 | B-Tier 1B | 399 | 854 | 384 | 460 | 853 | 680 |
| SS1196.1 | B-Tier 1B | 148 | 242 | 223 | 88 | 114 | 304 |
| 6535.3 | B-Tier 1B | 21 | 107 | 33 | 81 | 177 | 1518 |
| QH0692.42 | B-Tier 2 | 56 | 49 | 89 | 61 | 27 | 57 |
| SC422661.8 | B-Tier 2 | 73 | 220 | 97 | 68 | 37 | 65 |
| TRO.11 | B-Tier 2 | 60 | 68 | 55 | 68 | 41 | 187 |
| AC10.0.29 | B-Tier 2 | 21 | <20 | 109 | 34 | <20 | 241 |
| RHPA4259.7 | B-Tier 2 | 104 | 162 | 96 | 32 | 87 | 52 |
| THRO4156.18 | B-Tier 2 | 40 | 32 | 26 | 51 | 38 | <20 |
| REJO4541.67 | B-Tier 2 | 123 | 92 | <20 | 312 | 34 | 23 |
| WITO4160.33 | B-Tier 2 | 54 | 102 | 47 | 31 | 23 | 156 |
| CAAN5342.A2 | B-Tier 2 | 36 | 31 | 70 | 38 | 73 | 256 |
| PVO.4 | B-Tier 3 | 85 | 113 | 214 | 134 | 23 | 25 |
| TRJO4551.58 | B-Tier 3 | 43 | 71 | 21 | <20 | 46 | <20 |
| Neg. Control | MuLV | <20 | <20 | <20 | <20 | <20 | <20 |

| Breadth | ID80 titers |
|---|---|
| >90% | 1000 - 10000 |
| 70-90% | 250 - 1000 |
| 50-70% | 50 - 250 |
| 20-50% | 20 - 50 |
| 0 - 20% | <20 |

### Table S2. Sequence analysis of BG24 family antibodies isolated from donor 391370.

| | Heavy chain | | | | | | Light chain | | | | | |
|---|---|---|---|---|---|---|---|---|---|---|---|---|
| **mAb name** | **V-gene** | **% nt mutation** | **% AA mutation** | **CDRH3 (AA)** | **CDRH3 (seq)** | **Special features** | **V-gene** | **% nt mutation** | **% AA mutation** | **CDRL3 (AA)** | **CDRL3 (seq)** | **Special features** |
| BG3 | IGHV1-2*02 | 18.0% | 31.3% | 16 | ATRVRLPSSDDYGFDV | 6-nt insertion FWR3, 3-nt FWR1 deletion | IGLV2-11*01 | 7.1% | 14.6% | 5 | SSYEF | 12-nt deletion CDR1 |
| BG4 | IGHV1-2*02 | 20.8% | 34.4% | 16 | ATRVKLPSSDDYGFDV | 6-nt insertion FWR3, 3-nt FWR1 deletion | IGLV2-11*01 | 7.8% | 14.6% | 5 | SSYEF | 12-nt deletion CDR1 |
| BG5 | IGHV1-2*02 | 17.6% | 32.3% | 16 | ATRVKLPSSDDYGFDV | 6-nt insertion FWR3, 3-nt FWR1 deletion | IGLV2-11*01 | 7.1% | 13.5% | 5 | SSYEF | 12-nt deletion CDR1 |
| BG6 | IGHV1-2*02 | 20.1% | 34.4% | 16 | ATRVKLPSSDDYGFDV | 6-nt insertion FWR3, 3-nt FWR1 deletion | IGLV2-11*01 | 7.5% | 14.6% | 5 | SSYEF | 12-nt deletion CDR1 |
| BG7 | IGHV1-2*02 | 17.3% | 31.3% | 16 | ATRVKLPSSDDYGFDV | 6-nt insertion FWR3, 3-nt FWR1 deletion | IGLV2-11*01 | 7.1% | 14.6% | 5 | SSYEF | 12-nt deletion CDR1 |
| BG8 | IGHV1-2*02 | 19.0% | 32.3% | 17 | ATRVKVAPSSDSYGFDV | 6-nt insertion FWR3, 3-nt FWR1 deletion | IGLV2-11*01 | 8.6% | 16.9% | 5 | SSYEF | 12-nt deletion CDR1 |
| BG9 | IGHV1-2*02 | 19.0% | 30.6% | 16 | ARQTLVTASTSFAFNN | 3-nt insertion CDR1 | IGLV2-11*01 | 5.8% | 12.9% | 5 | SSYEY | 21-nt deletion CDR1 |
| BG10 | IGHV1-2*02 | 16.0% | 25.5% | 16 | ARQTLRSASASFAFNS | 6-nt insertion CDR1 | IGLV2-11*01 | 10.1% | 14.1% | 5 | SSNEF | 21-nt deletion CDR1 |
| BG22 | IGHV1-2*02 | 13.7% | 23.7% | 16 | ASNVKVSSSTQYVFDV | 6-nt insertion FWR3 | IGLV2-11*01 | 9.5% | 16.1% | 5 | SAFEF | 18-nt deletion CDR1 |
| BG23 | IGHV1-2*02 | 11.4% | 21.9% | 16 | ATQVKLSSSIEYPFNI | 3-nt FWR1 deletion | IGLV2-11*01 | 7.3% | 16.1% | 5 | SAFEF | 18-nt deletion CDR1 |
| **BG24** | **IGHV1-2*02** | **13.4%** | **22.7%** | **16** | **ATQVKLDSSAGYPFDI** | **-** | **IGLV2-11*01** | **8.0%** | **19.5%** | **5** | **SAFEY** | **18-nt deletion CDR1** |
| BG29 | IGHV1-2*02 | 19.4% | 30.6% | 16 | ARQTLVTASTSFAFNN | 3-nt insertion CDR1 | IGLV2-11*01 | 5.8% | 12.9% | 5 | SSYEY | 21-nt deletion CDR1 |
| BG30 | IGHV1-2*02 | 15.8% | 24.7% | 16 | ATRVKVTSSTTFTFDI | - | IGLV2-11*01 | 9.2% | 17.2% | 5 | SAFEY | 18-nt deletion CDR1 |
| BG31 | IGHV1-2*02 | 19.0% | 34.4% | 16 | ATRVKLPSSDDYGFDV | 6-nt insertion FWR3, 3-nt FWR1 deletion | IGLV2-11*01 | 7.8% | 15.7% | 5 | SSYEF | 12-nt deletion CDR1 |
| BG32 | IGHV1-2*02 | 17.6% | 30.2% | 16 | ATRVKLPSSDDYGFDV | 6-nt insertion FWR3, 3-nt FWR1 deletion | IGLV2-11*01 | 7.8% | 16.9% | 5 | SSYEF | 12-nt deletion CDR1 |
| BG33 | IGHV1-2*02 | 14.7% | 26.8% | 16 | ASQVAVGDSTHYPFDV | - | IGLV2-11*01 | 6.9% | 12.8% | 5 | SAFEF | 18-nt deletion CDR1 |
| BG34 | IGHV1-2*02 | 17.0% | 31.3% | 16 | ATRVKLPSSDDYGFDV | 6-nt insertion FWR3, 3-nt FWR1 deletion | IGLV2-11*01 | 7.1% | 14.6% | 5 | SSYEF | 12-nt deletion CDR1 |
| BG35 | IGHV1-2*02 | 16.2% | 26.8% | 16 | ASNVKVPSSTVYVFDV | 6-nt insertion FWR3 | IGLV2-11*01 | 8.8% | 16.1% | 5 | SAFEF | 18-nt deletion CDR1 |
| BG36 | IGHV1-2*02 | 19.0% | 30.6% | 16 | ARQTLVTASTSFAFNN | 3-nt insertion CDR1 | IGLV2-11*01 | 5.8% | 12.9% | 5 | SSYEY | 21-nt deletion CDR1 |
| BG37 | IGHV1-2*02 | 17.1% | 25.8% | 18 | ATRVRLPSSSSGDYAFNI | - | IGLV2-11*01 | 8.8% | 18.4% | 5 | SSYEF | 18-nt deletion CDR1 |
| BG38 | IGHV1-2*02 | 18.2% | 28.9% | 16 | ATQVAVAASTGWAFDV | 6-nt insertion FWR3 | IGLV2-11*01 | 10.7% | 17.2% | 5 | SAFEY | 18-nt deletion CDR1 |
| BG39 | IGHV1-2*02 | 14.1% | 24.7% | 16 | ASNVKVTSSTQFVFDV | 6-nt insertion FWR3 | IGLV2-11*01 | 9.2% | 16.1% | 5 | SAFEF | 18-nt deletion CDR1 |
| BG40 | IGHV1-2*02 | 16.6% | 32.3% | 16 | AVSTLMASSSTWGLNV | 6-nt insertion FWR3 | IGLV2-11*01 | 9.5% | 18.4% | 5 | SSYEF | 18-nt deletion CDR1 |
| BG41 | IGHV1-2*02 | 13.8% | 29.2% | 12 | AVSTLSDFGFNI | 6-nt insertion FWR3 | IGLV2-11*01 | 9.2% | 12.6% | 5 | SSYEF | 18-nt deletion CDR1 |
| BG42 | IGHV1-2*02 | 13.4% | 23.7% | 16 | ASNVKVPSSTQFVFDV | 6-nt insertion FWR3 | IGLV2-11*01 | 9.2% | 17.2% | 5 | SAFEF | 18-nt deletion CDR1 |
| BG46 | IGHV1-2*02 | 20.1% | 34.4% | 16 | ATRVKLPSSDDYGFDV | 6-nt insertion FWR3, 3-nt FWR1 deletion | IGLV2-11*01 | 7.5% | 14.6% | 5 | SSYEF | 12-nt deletion CDR1 |
| BG47 | IGHV1-2*02 | 13.6% | 24.2% | 16 | ATQVKLSSSIEYPFDI | 3-nt FWR1 deletion | IGLV2-11*01 | 6.9% | 16.1% | 5 | SAFEF | 18-nt deletion CDR1 |
| BG48 | IGHV1-2*02 | 19.0% | 29.6% | 16 | ARQTLVTASTSFAFNN | 3-nt insertion CDR1 | IGLV2-11*01 | 5.8% | 12.9% | 5 | SSYEY | 21-nt deletion CDR1 |
| BG49 | IGHV1-2*02 | 19.0% | 30.6% | 16 | ARQTLVTASTSFAFNN | 3-nt insertion CDR1 | IGLV2-11*01 | 5.8% | 12.9% | 5 | SSYEY | 21-nt deletion CDR1 |
| BG50 | IGHV1-2*02 | 13.4% | 23.7% | 16 | ASNVKVPSSTQFVFDV | 6-nt insertion FWR3 | IGLV2-11*01 | 9.2% | 17.2% | 5 | SAFEF | 18-nt deletion CDR1 |
| BG51 | IGHV1-2*02 | 16.6% | 30.2% | 16 | ATRVKLPSSDDYGFDV | 6-nt insertion FWR3, 3-nt FWR1 deletion | IGLV2-11*01 | 7.1% | 14.6% | 5 | SSYEF | 12-nt deletion CDR1 |
| BG52 | IGHV1-2*02 | 16.2% | 31.3% | 16 | AVKVLRTSSSGYGFNV | 6-nt insertion FWR3 | IGLV2-11*01 | 6.5% | 12.6% | 5 | SSYEY | 18-nt deletion CDR1 |
| BG53 | IGHV1-2*02 | 12.1% | 24.0% | 16 | ATQVKLSSSIEYPFNI | 3-nt FWR1 deletion | IGLV2-11*01 | 6.9% | 14.9% | 5 | SAFEF | 18-nt deletion CDR1 |
| BG54 | IGHV1-2*02 | 13.7% | 23.7% | 16 | ASNVKVSSSTQYVFDV | 6-nt insertion FWR3 | IGLV2-11*01 | 9.5% | 16.1% | 5 | SAFEF | 18-nt deletion CDR1 |
| BG55 | IGHV1-2*02 | 19.0% | 30.6% | 16 | ARQTLVTASTSFAFNN | 3-nt insertion CDR1 | IGLV2-11*01 | 5.8% | 12.9% | 5 | SSYEY | 21-nt deletion CDR1 |
| BG56 | IGHV1-2*02 | 19.0% | 30.6% | 16 | ARQTLVTASTSFAFNN | 3-nt insertion CDR1 | IGLV2-11*01 | 5.8% | 12.9% | 5 | SSYEY | 21-nt deletion CDR1 |
| BG57 | IGHV1-2*02 | 19.0% | 33.7% | 16 | ARQTLTTASTSFAFNH | - | IGLV2-11*01 | 7.0% | 14.1% | 5 | SSYEF | 21-nt deletion CDR1 |
| BG58 | IGHV1-2*02 | 16.2% | 31.3% | 16 | AVKVLRTSSSGYGFNV | 6-nt insertion FWR3 | IGLV2-11*01 | 6.5% | 12.6% | 5 | SSYEY | 18-nt deletion CDR1 |
| BG59 | IGHV1-2*02 | 17.0% | 29.6% | 16 | ARQTLVTASTSFAFDS | - | IGLV2-11*01 | 5.5% | 11.9% | 5 | SSLEF | 21-nt deletion CDR1 |

### Table S3. 126 virus cross-clade neutralization panel data.

| Virus ID | Clade* | BG24 WT | | BG24 G54W | | BG24 CDR2-v1 | | BG24 CDR2-v2 | | BG24 $Y100_BW$ | |
|---|---|---|---|---|---|---|---|---|---|---|---|
| | | IC50 | IC80 | IC50 | IC80 | IC50 | IC80 | IC50 | IC80 | IC50 | IC80 |
| 6535.3 | B | 1.55 | 5.26 | 1.57 | 17.7 | 5.55 | >25 | 6.58 | >25 | >25 | >25 |
| QH0692.42 | B | 0.96 | 2.71 | 0.54 | 1.98 | 0.85 | 3.13 | 0.94 | 3.54 | 0.67 | 2.47 |
| SC422661.8 | B | 0.06 | 0.27 | 0.05 | 0.21 | 0.03 | 0.15 | 0.05 | 0.24 | 0.08 | 0.30 |
| PVO.4 | B | 0.18 | 0.83 | 0.10 | 0.67 | 0.17 | 1.17 | 0.21 | 0.96 | 0.14 | 0.93 |
| TRO.11 | B | 0.16 | 1.08 | 0.07 | 0.33 | 0.05 | 0.25 | 0.05 | 0.19 | 0.13 | 0.72 |
| AC10.0.29 | B | 5.21 | >25 | 7.27 | >25 | 19.96 | >25 | 11.20 | >25 | >25 | >25 |
| RHPA4259.7 | B | 0.03 | 0.09 | 0.02 | 0.08 | 0.01 | 0.05 | 0.02 | 0.05 | 0.02 | 0.08 |
| THRO4156.18 | B | 3.01 | 18.7 | 1.20 | 6.04 | 0.98 | 5.33 | 1.20 | 6.38 | 4.48 | 18.0 |
| REJO4541.67 | B | 0.10 | 0.45 | 0.16 | 1.39 | 0.12 | 0.93 | 0.13 | 0.88 | >25 | >25 |
| TRJO4551.58 | B | 0.34 | 1.69 | 0.83 | 4.87 | 0.11 | 0.53 | 0.19 | 0.69 | >25 | >25 |
| WITO4160.33 | B | 0.15 | 0.49 | 0.10 | 0.36 | 0.09 | 0.34 | 0.11 | 0.36 | 0.13 | 0.63 |
| CAAN5342.A2 | B | 1.08 | 3.96 | 0.83 | 3.05 | 1.24 | 5.95 | 0.71 | 2.44 | 0.85 | 4.42 |
| WEAU_d15_410_787 | B (T/F) | 0.07 | 0.32 | 0.06 | 0.19 | 0.06 | 0.21 | 0.04 | 0.13 | 0.13 | 0.67 |
| 1006_11_C3_1601 | B (T/F) | 0.08 | 0.33 | 0.07 | 0.24 | 0.09 | 0.30 | 0.12 | 0.39 | 0.09 | 0.33 |
| 1054_07_TC4_1499 | B (T/F) | 0.69 | 3.42 | 0.32 | 1.77 | 0.50 | 2.99 | 0.48 | 2.44 | 0.83 | 5.03 |
| 1056_10_TA11_1826 | B (T/F) | 0.39 | 1.88 | 0.16 | 1.23 | 0.20 | 1.43 | 0.34 | 1.50 | 0.55 | 2.70 |
| 1012_11_TC21_3257 | B (T/F) | 0.11 | 0.50 | 0.05 | 0.21 | 0.08 | 0.46 | 0.06 | 0.28 | 0.09 | 0.41 |
| 6240_08_TA5_4622 | B (T/F) | 0.41 | 2.82 | 0.35 | 1.62 | 0.38 | 1.75 | 0.37 | 1.67 | 0.54 | 2.60 |
| 6244_13_B5_4576 | B (T/F) | 0.22 | 0.76 | 0.15 | 0.52 | 0.17 | 0.56 | 0.16 | 0.53 | 0.16 | 0.57 |
| 62357_14_D3_4589 | B (T/F) | 0.38 | 3.74 | 0.13 | 0.96 | 0.09 | 0.40 | 0.17 | 0.77 | >25 | >25 |
| SC05_8C11_2344 | B (T/F) | 0.79 | 2.63 | 0.61 | 2.05 | 0.70 | 2.44 | 0.50 | 1.39 | 15.47 | >25 |
| Du156.12 | C | 0.07 | 0.35 | 0.03 | 0.14 | 0.02 | 0.09 | 0.02 | 0.06 | 0.20 | 1.34 |
| Du172.17 | C | 24.66 | >25 | 0.46 | 5.08 | 0.05 | 0.19 | 0.12 | 1.07 | >25 | >25 |
| Du422.1 | C | 8.38 | >25 | 1.82 | 7.22 | 0.36 | 1.63 | 0.09 | 0.31 | >25 | >25 |
| ZM197M.PB7 | C | 0.36 | 1.25 | 0.19 | 0.91 | 0.14 | 0.50 | 0.17 | 0.83 | 0.30 | 1.51 |
| ZM214M.PL15 | C | 0.40 | 2.71 | 0.12 | 0.76 | 0.11 | 1.10 | 0.15 | 0.97 | 2.38 | 17.7 |
| ZM233M.PB6 | C | 6.55 | >25 | 0.67 | 2.92 | 0.44 | 1.87 | 0.21 | 0.68 | >25 | >25 |
| ZM249M.PL1 | C | 0.12 | 0.40 | 0.05 | 0.18 | 0.04 | 0.12 | 0.03 | 0.11 | 0.31 | 2.33 |
| ZM53M.PB12 | C | 0.33 | 1.51 | 0.25 | 0.83 | 0.27 | 1.30 | 0.26 | 1.18 | 1.00 | 6.29 |
| ZM109F.PB4 | C | 0.31 | 2.13 | 0.08 | 0.41 | 0.05 | 0.16 | 0.03 | 0.10 | >25 | >25 |
| ZM135M.PL10a | C | 0.13 | 0.48 | 0.13 | 0.54 | 0.13 | 0.60 | 0.12 | 0.80 | 0.63 | 7.58 |
| CAP45.2.00.G3 | C | 0.04 | 0.25 | 0.02 | 0.11 | 0.01 | 0.03 | 0.02 | 0.05 | 3.72 | >25 |
| CAP210.2.00.E8 | C | >25 | >25 | 6.96 | >25 | 9.99 | >25 | 2.43 | 18.72 | >25 | >25 |
| HIV-001428-2.42 | C | 0.03 | 0.09 | 0.02 | 0.06 | 0.01 | 0.05 | 0.02 | 0.08 | 2.48 | >25 |
| HIV-0013095-2.11 | C | 1.29 | 15.4 | 1.28 | 8.92 | 1.69 | >25 | 2.86 | >25 | >25 | >25 |
| HIV-16055-2.3 | C | 0.08 | 0.31 | 0.05 | 0.14 | 0.04 | 0.13 | 0.04 | 0.10 | 0.11 | 0.35 |
| HIV-16845-2.22 | C | 3.52 | 24.8 | 1.92 | 8.97 | 1.97 | 15.8 | 1.71 | 8.63 | 4.00 | >25 |
| Ce1086_B2 | C (T/F) | 0.39 | 1.02 | 0.16 | 0.68 | 0.18 | 0.60 | 0.18 | 0.56 | 1.13 | 13.1 |
| Ce0393_C3 | C (T/F) | 0.10 | 0.45 | 0.06 | 0.29 | 0.04 | 0.15 | 0.05 | 0.19 | 0.37 | 2.00 |
| Ce1176_A3 | C (T/F) | 0.89 | 4.41 | 0.40 | 1.88 | 0.55 | 2.56 | 0.58 | 2.56 | 6.61 | >25 |
| Ce2010_F5 | C (T/F) | 0.25 | 1.24 | 0.12 | 0.56 | 0.17 | 0.78 | 0.31 | 1.06 | 0.25 | 1.14 |
| Ce0682_E4 | C (T/F) | 0.05 | 0.26 | 0.03 | 0.08 | 0.02 | 0.08 | 0.04 | 0.15 | 0.05 | 0.21 |
| Ce1172_H1 | C (T/F) | >25 | >25 | >25 | >25 | >25 | >25 | >25 | >25 | >25 | >25 |
| Ce2060_G9 | C (T/F) | 0.23 | 1.10 | 0.13 | 0.60 | 0.17 | 0.78 | 0.12 | 0.53 | 0.33 | 2.36 |
| Ce703010054_2A2 | C (T/F) | 0.69 | 2.47 | 0.17 | 0.83 | 0.07 | 0.32 | 0.09 | 0.40 | >25 | >25 |
| BF1266.431a | C (T/F) | >25 | >25 | 5.23 | >25 | 8.31 | >25 | >25 | >25 | >25 | >25 |
| 246F C1G | C (T/F) | >25 | >25 | >25 | >25 | 4.30 | >25 | 11.08 | >25 | >25 | >25 |
| 249M B10 | C (T/F) | 0.35 | 1.60 | 0.13 | 0.60 | 0.12 | 0.44 | 0.18 | 0.60 | 1.48 | 8.16 |
| ZM247v1(Rev-) | C (T/F) | 0.11 | 0.57 | 0.09 | 0.45 | 0.09 | 0.43 | 0.08 | 0.55 | 0.60 | 3.53 |
| 7030102001E5(Rev-) | C (T/F) | 2.05 | 7.09 | 1.67 | 5.45 | 0.84 | 2.67 | 0.70 | 2.14 | >25 | >25 |
| 1394C9G1(Rev-) | C (T/F) | 0.53 | 2.49 | 0.17 | 1.12 | 0.25 | 1.78 | 0.22 | 1.00 | 6.44 | >25 |
| Ce704809221_1B3 | C (T/F) | 0.78 | 3.69 | 0.15 | 1.06 | 0.13 | 0.95 | 0.19 | 1.18 | 17.33 | >25 |
| CNE19 | BC | 0.06 | 0.28 | 0.04 | 0.17 | 0.04 | 0.21 | 0.04 | 0.22 | 0.06 | 0.36 |
| CNE20 | BC | 0.47 | 2.45 | 0.19 | 1.02 | 0.19 | 0.92 | 0.32 | 2.72 | >25 | >25 |
| CNE21 | BC | 0.60 | 2.77 | 0.20 | 1.42 | 0.22 | 0.99 | 0.15 | 0.88 | >25 | >25 |
| CNE17 | BC | 1.00 | 4.88 | 0.59 | 2.76 | 0.79 | 3.60 | 0.96 | 4.69 | >25 | >25 |
| CNE30 | BC | 0.87 | 2.90 | 0.55 | 1.75 | 0.41 | 1.35 | 0.34 | 1.13 | 2.88 | 16.6 |
| CNE52 | BC | 0.07 | 0.22 | 0.03 | 0.08 | 0.02 | 0.06 | 0.02 | 0.07 | 0.10 | 0.40 |
| CNE53 | BC | 0.16 | 0.60 | 0.10 | 0.33 | 0.09 | 0.31 | 0.09 | 0.39 | 0.19 | 0.98 |
| CNE58 | BC | 0.04 | 0.10 | 0.03 | 0.08 | 0.02 | 0.06 | 0.03 | 0.07 | 0.05 | 0.12 |
| MS208.A1 | A | 0.32 | 1.95 | 0.10 | 0.32 | 0.07 | 0.22 | 0.06 | 0.25 | 7.33 | >25 |
| Q23.17 | A | 0.10 | 0.40 | 0.05 | 0.15 | 0.04 | 0.14 | 0.04 | 0.16 | 0.12 | 0.77 |
| Q461.e2 | A | 0.48 | 1.54 | 0.27 | 0.69 | 0.28 | 0.74 | 0.18 | 0.61 | 0.27 | 0.98 |
| Q769.d22 | A | 0.09 | 0.36 | 0.05 | 0.17 | 0.05 | 0.16 | 0.06 | 0.19 | 0.09 | 0.33 |
| Q259.d2.17 | A | >25 | >25 | >25 | >25 | 1.02 | 12.3 | 0.09 | 0.30 | >25 | >25 |
| Q842.d12 | A | 0.05 | 0.14 | 0.03 | 0.08 | 0.03 | 0.09 | 0.03 | 0.08 | 0.02 | 0.09 |
| 0260.v5.c36 | A | 0.54 | 1.75 | 0.35 | 1.11 | 0.32 | 1.03 | 0.26 | 0.83 | 0.48 | 2.21 |
| 3415.v1.c1 | A | 0.29 | 0.93 | 0.12 | 0.38 | 0.08 | 0.27 | 0.06 | 0.20 | 6.65 | >25 |
| 3365.v2.c2 | A | 0.06 | 0.18 | 0.04 | 0.09 | 0.04 | 0.12 | 0.04 | 0.12 | 0.08 | 0.36 |
| 191955_A11 | A (T/F) | >25 | >25 | >25 | >25 | >25 | >25 | 11.5 | >25 | >25 | >25 |
| 191084 B7-19 | A (T/F) | 0.17 | 0.37 | 0.11 | 0.31 | 0.08 | 0.18 | 0.11 | 0.29 | 0.16 | 0.60 |
| 9004SS_A3_4 | A (T/F) | 0.42 | 1.98 | 0.25 | 1.72 | 0.15 | 0.69 | 0.23 | 0.97 | 1.33 | 8.80 |
| T257-31 | AG | 1.41 | 11.9 | 0.70 | 5.42 | 0.72 | 3.50 | 0.49 | 2.26 | 2.48 | 13.9 |
| 928-28 | AG | 0.55 | 1.79 | 0.31 | 1.30 | 0.29 | 0.94 | 0.33 | 1.44 | 1.95 | 8.13 |
| 263-8 | AG | 0.14 | 0.41 | 0.05 | 0.19 | 0.06 | 0.22 | 0.07 | 0.47 | 0.13 | 1.05 |
| T250-4 | AG | >25 | >25 | >25 | >25 | >25 | >25 | >25 | >25 | >25 | >25 |
| T251-18 | AG | 0.92 | 4.68 | 0.45 | 1.63 | 0.37 | 1.70 | 0.40 | 1.44 | 3.29 | 17.7 |
| T278-50 | AG | >25 | >25 | >25 | >25 | >25 | >25 | >25 | >25 | >25 | >25 |
| T255-34 | AG | 0.16 | 0.89 | 0.05 | 0.26 | 0.02 | 0.13 | 0.03 | 0.16 | 2.08 | >25 |
| 211-9 | AG | >25 | >25 | >25 | >25 | 0.97 | 5.11 | 1.02 | 4.80 | >25 | >25 |
| 235-47 | AG | 0.40 | 3.30 | 0.07 | 0.34 | 0.03 | 0.10 | 0.03 | 0.10 | >25 | >25 |
| 620345.c01 | AE | >25 | >25 | >25 | >25 | >25 | >25 | >25 | >25 | >25 | >25 |
| CNE8 | AE | 0.28 | 1.21 | 0.12 | 0.54 | 0.07 | 0.32 | 0.10 | 0.44 | 1.49 | 15.0 |
| C1080.c03 | AE | 1.27 | 6.28 | 0.39 | 2.69 | 0.25 | 1.67 | 0.30 | 1.92 | 14.77 | >25 |
| R2184.c04 | AE | 0.07 | 0.29 | 0.02 | 0.10 | 0.03 | 0.11 | 0.03 | 0.14 | 0.07 | 0.29 |
| R1166.c01 | AE | 0.20 | 0.69 | 0.13 | 0.43 | 0.09 | 0.30 | 0.12 | 0.38 | 0.32 | 1.11 |
| R3265.c06 | AE | 0.51 | 4.70 | 0.06 | 1.18 | 0.17 | 1.22 | 0.14 | 0.86 | 8.17 | >25 |
| C2101.c01 | AE | 0.09 | 0.42 | 0.02 | 0.10 | 0.04 | 0.13 | 0.03 | 0.13 | 0.24 | 2.05 |
| C3347.c11 | AE | 0.04 | 0.18 | 0.01 | 0.05 | 0.01 | 0.04 | 0.02 | 0.07 | 0.17 | 1.93 |
| C4118.c09 | AE | 0.12 | 0.68 | 0.04 | 0.21 | 0.04 | 0.24 | 0.07 | 0.41 | 3.00 | >25 |
| CNE5 | AE | 0.96 | 5.22 | 0.18 | 0.80 | 0.11 | 0.48 | 0.11 | 0.49 | >25 | >25 |
| BJOX009000.02.4 | AE | 2.52 | 7.22 | 1.56 | 5.49 | 2.04 | 7.11 | 1.54 | 5.34 | 6.57 | >25 |
| BJOX015000.11.5 | AE (T/F) | 0.14 | 1.01 | 0.06 | 0.39 | 0.15 | 0.98 | 0.08 | 0.63 | 0.13 | 1.00 |
| BJOX010000.06.2 | AE (T/F) | >25 | >25 | >25 | >25 | >25 | >25 | 19.08 | >25 | >25 | >25 |
| BJOX025000.01.1 | AE (T/F) | >25 | >25 | 4.13 | >25 | 0.04 | 0.23 | 0.12 | 0.71 | >25 | >25 |
| BJOX028000.10.3 | AE (T/F) | 0.42 | 22.8 | 0.01 | 0.04 | 0.01 | 0.03 | 0.00 | 0.02 | >25 | >25 |
| X1193_c1 | G | 0.29 | 1.34 | 0.08 | 0.27 | 0.04 | 0.19 | 0.08 | 0.32 | >25 | >25 |
| P0402_c2_11 | G | 0.17 | 0.60 | 0.06 | 0.30 | 0.04 | 0.16 | 0.05 | 0.16 | 1.95 | 14.8 |
| X1254_c3 | G | 0.19 | 0.55 | 0.10 | 0.34 | 0.08 | 0.28 | 0.09 | 0.24 | 3.76 | >25 |
| X2088_c9 | G | >25 | >25 | >25 | >25 | >25 | >25 | >25 | >25 | >25 | >25 |
| X2131_C1_B5 | G | 1.05 | 3.29 | 0.19 | 1.56 | 0.08 | 0.50 | 0.12 | 0.76 | >25 | >25 |
| P1981_C5_3 | G | 7.34 | >25 | 1.63 | 6.44 | 0.48 | 1.67 | 0.44 | 1.51 | >25 | >25 |
| X1632_S2_B10 | G | >25 | >25 | >25 | >25 | 0.92 | >25 | 0.20 | >25 | >25 | >25 |
| 3016.v5.c45 | D | >25 | >25 | 23.60 | >25 | 2.29 | 20.3 | 1.39 | 16.7 | >25 | >25 |
| A07412M1.vrc12 | D | 0.15 | 0.57 | 0.10 | 0.35 | 0.08 | 0.39 | 0.08 | 0.35 | 0.23 | 1.23 |
| 231965.c01 | D | 1.23 | 3.52 | 0.86 | 1.53 | 0.78 | 1.38 | 0.57 | 1.23 | 3.44 | 13.6 |
| 231966.c02 | D | 0.35 | 1.66 | 0.15 | 0.73 | 0.08 | 0.28 | 0.09 | 0.34 | 23.80 | >25 |
| 6405.v4.c34 | D | 0.91 | 1.97 | 0.50 | 1.59 | 0.40 | 1.03 | 0.26 | 0.83 | 0.67 | 2.26 |
| 3817.v2.c59 | CD | >25 | >25 | >25 | >25 | 14.01 | >25 | 21.18 | >25 | >25 | >25 |
| 6480.v4.c25 | CD | 0.01 | 0.06 | 0.01 | 0.04 | 0.01 | 0.03 | 0.01 | 0.03 | 0.02 | 0.08 |
| 6952.v1.c20 | CD | 0.09 | 0.42 | 0.05 | 0.18 | 0.03 | 0.17 | 0.04 | 0.18 | 0.14 | 1.11 |
| 6811.v7.c18 | CD | 0.13 | 0.63 | 0.08 | 0.36 | 0.08 | 0.30 | 0.06 | 0.22 | 0.08 | 0.63 |
| 89-F1_2_25 | CD | >25 | >25 | >25 | >25 | >25 | >25 | >25 | >25 | >25 | >25 |
| 3301.v1.c24 | AC | 0.04 | 0.16 | 0.03 | 0.11 | 0.03 | 0.11 | 0.04 | 0.13 | 0.04 | 0.17 |
| 6041.v3.c23 | AC | 0.05 | 0.27 | 0.01 | 0.08 | 0.01 | 0.03 | 0.01 | 0.02 | 2.33 | >25 |
| 6540.v4.c1 | AC | >25 | >25 | >25 | >25 | >25 | >25 | >25 | >25 | >25 | >25 |
| 6545.v4.c1 | AC | >25 | >25 | >25 | >25 | >25 | >25 | >25 | >25 | >25 | >25 |
| 0815.v3.c3 | ACD | 0.03 | 0.13 | 0.01 | 0.04 | 0.01 | 0.04 | 0.02 | 0.05 | 0.03 | 0.13 |
| 3103.v3.c10 | ACD | 1.52 | 4.13 | 0.86 | 2.68 | 0.17 | 0.54 | 0.23 | 0.59 | >25 | >25 |
| Ce703010217_B6 | C | 0.13 | 0.66 | 0.07 | 0.25 | 0.05 | 0.25 | 0.11 | 0.39 | 0.19 | 0.73 |
| CNE55 | AE | 0.24 | 1.09 | 0.11 | 0.51 | 0.05 | 0.22 | 0.10 | 0.45 | >25 | >25 |
| 246-F3_C10_2 | AC | 0.17 | 0.57 | 0.11 | 0.48 | 0.07 | 0.19 | 0.09 | 0.32 | 0.36 | 1.93 |
| CH119.10 | CRF07 | 0.28 | 1.27 | 0.16 | 0.54 | 0.15 | 0.67 | 0.13 | 0.59 | 0.28 | 1.58 |
| BJOX002000.03.2 | CRF07 | 0.34 | 1.84 | 0.13 | 0.47 | 0.09 | 0.41 | 0.09 | 0.45 | 8.07 | >25 |
| 25710-2.43 | C | 0.23 | 1.54 | 0.17 | 0.86 | 0.10 | 0.72 | 0.13 | 0.66 | 0.49 | 3.63 |
| AD8.MM | B | 0.31 | 0.88 | 0.13 | 0.36 | 0.21 | 0.49 | 0.25 | 0.70 | 0.19 | 0.71 |
| MuLV | Neg. Con. | >25 | >25 | >25 | >25 | >25 | >25 | >25 | >25 | >25 | >25 |

| IC50 (μg/mL) | |
|---|---|
| >25 | white |
| 1.0 - 25 | yellow |
| 0.1 - 1.0 | orange |
| <0.1 | red |

BG24 – $IC_{50}$=0.29μg/mL, 84.9% breadth; $BG24_{G54W}$ – $IC_{50}$=0.15μg/mL, 88.1% breadth; $BG24_{PG20\text{-}CDR2\text{-}v1}$ – $IC_{50}$=0.15μg/mL, 92.9% breadth; $BG24_{PG20\text{-}CDR2\text{-}v2}$ – $IC_{50}$=0.14μg/mL, 92.1% breadth; BG24 $Y100_DW$ – IC50=0.52μg/mL, 66.7% breadth

**Table S4. X-ray data collection, refinement and validation statistics.**

| | **BG24 Fab** *(12-2, SSRL)* | **BG24-BG505-101074** *(12-2, SSRL)* |
|---|---|---|
| **PDB ID** | | |
| **Data collection**[a] | | |
| Space group | $C222_1$ | H3 |
| Unit cell (Å) | 66.5, 120, 120 | 209, 209, 156 |
| α, β, γ (°) | 90, 90, 90 | 90, 90, 120 |
| Wavelength (Å) | 1.0 | 1.0 |
| Resolution (Å) | 39.28-2.0 (2.06-2.0) | 39.28-3.8 (4.06-3.80) |
| Unique Reflections | 33,354 (2,031) | 24,941 (4,480) |
| Completeness (%) | 99 (94.2) | 99.6 (99.2) |
| Redundancy | 45.1 (30.1) | 11.3 (11.2) |
| $CC_{1/2}$ (%) | 95.9 (94.2) | 99.8 (21.0) |
| <I/σI> | 11.6 (0.8) | 9.7 (0.9) |
| Mosaicity (˚) | 0.20 | 0.26 |
| $R_{merge}$ (%) | 22.7 (389) | 14.2 (406) |
| $R_{pim}$ (%) | 3.8 (77.9) | 3.7 (109) |
| Wilson *B* -factor | 33.9 | 188.0 |
| **Refinement and Validation** | | |
| Resolution (Å) | 34.2-2.1 | 39.2-3.8 |
| Number of atoms | | |
| Protein | 3,208 | 11,159 |
| Ligand | 27 | 792 |
| Water | 52 | 0 |
| $R_{work}/R_{free}$ (%) | 20.4/24.5 | 25.7/27.9 |
| R.m.s. deviations | | |
| Bond lengths (Å) | 0.004 | 0.0055 |
| Bond angles (˚) | 0.716 | 0.82 |
| MolProbity score | 2.54 | 1.63 |
| Clashscore (all atom) | 5.9 | 6.2 |
| Poor rotamers (%) | 5 | 0 |
| Ramachandran plot | | |
| Favored (%) | 96 | 92.9 |
| Allowed (%) | 3.5 | 6.2 |
| Disallowed (%) | 0.5 | 0.9 |
| Average *B* -factor (Å) | 68.1 | 262.6 |

[a]Numbers in parentheses correspond to the highest resolution shell

**Table S5. Cryo-EM data collection, refinement and validation statistics.**

| | $BG24_{CDR\text{-}v2}$ DU422 SOSIP.664 v4.2 10-1074 |
|---|---|
| **PDB** | |
| **EMDB** | |
| **Data collection and processing** | |
| Microscope | Talos Arctica |
| Camera | Gatan K3 DED |
| Magnification | 45,000x |
| Voltage (kV) | 200 |
| Recording mode | counting |
| Dose rate ($e^-$/pixel/s) | 13.5 |
| Electron dose ($e^-$/Å$^2$) | 60 |
| Defocus range (µm) | 1.0 - 2.5 |
| Pixel size (Å) | 0.869 |
| Micrographs collected | 1,989 |
| Micrographs used | 1,180 |
| Total extracted particles | 455,671 |
| Refined particles | 248,600 |
| **Reconstruction** | |
| Final particles | 204,220 |
| Symmetry imposed | C3 |
| Nominal Resolution (Å) | |
| FSC 0.143 (unmasked.masked) | 4.1/3.5 |
| Map sharpening *B* -factor | -110 |
| **Refinement and Validation** | |
| Number of atoms | |
| Protein | 20,391 |
| Ligand | 2,271 |
| MapCC (global/local) | 0.79/0.75 |
| R.m.s. deviations | |
| Bond lengths (Å) | 0.01 |
| Bond angles (˚) | 1.27 |
| MolProbity score | 1.78 |
| Clashscore (all atom) | 11.1 |
| Poor rotamers (%) | 0.1 |
| Ramachandran plot | |
| Favored (%) | 91.8 |
| Allowed (%) | 7.9 |
| Disallowed (%) | 0.3 |

**Table S6. Neutralization testing of BG24 and VRC01 on an extended HIV$_{YU2}$ pseudovirus site mutant panel**

| YU2 gp160 mutation | Region | Mutation | BG24 $IC_{50}$ (µg/ml) | VRC01 $IC_{50}$ (µg/ml) | BG24 Fold change relative to YU2 Wildtype | VRC01 Fold change relative to YU2 Wildtype |
|---|---|---|---|---|---|---|
| | | YU2 Wildtype | 0.05 | 0.06 | | |
| YU2 gp160 mutation | | E102K | 0.02 | 0.06 | | |
| | V2 Loop | N160K | 0.02 | 0.03 | | |
| | V2 Loop | N160T | 0.03 | 0.02 | | |
| | V2 Loop | T162I | 0.03 | 0.05 | | |
| | V2 Loop | T162N | 0.02 | 0.05 | | |
| | D loop | N276D | 0.02 | 0.01 | | |
| | D loop | T278K | <0,01 | 0.01 | | |
| | D loop | T278A | <0,01 | 0.01 | | |
| | D loop | T278I | <0,01 | <0.01 | | |
| | D loop | N279K | >25 | >25 | | |
| | D loop | N279H | >25 | 0.02 | | |
| | D loop | N280Y | >25 | >25 | | |
| | D loop | A281T | >25 | 11.95 | | |
| | D loop | T278I/A281T | >25 | >25 | | |
| | | N295S | 0.04 | 0.01 | | |
| | V3 Loop | N301D | <0,01 | <0.01 | | |
| | V3 Loop | N332K | 0.02 | 0.02 | | |
| | V3 Loop | N332S | 0.02 | 0.04 | | |
| | V3 Loop | N332Y | 0.02 | 0.04 | | |
| | V3 Loop | S334D | 0.04 | 0.05 | | |
| | V3 Loop | S334N | 0.02 | 0.02 | | |
| | CD4 binding loop | G366E | 0.01 | <0.01 | | |
| | CD4 binding loop | N365L | 0.01 | <0.01 | | |
| | CD4 binding loop | I371M | 0.03 | 0.02 | | |
| | | E429K | 0.04 | 0.06 | | |
| | | N448Q | 0.04 | 0.11 | | |
| | β23/V5 loop | G458D | 0.10 | 0.17 | | |
| | β23/V5 loop | G459D | 0.03 | 0.02 | | |
| | β23/V5 loop | G471R | 0.04 | 0.04 | | |

mAb titers (µg/ml)
- <0.01
- 0.01 - 0.1
- 0.1 - 1.0
- 1.0 - 20.0
- ND (>10/>20)

Fold change
- < 0.33
- 0.33 - 3
- 3-10
- > 10
